# Deep Cellular and Spatial Profiling of the Mouse Spinal Cord Reveals Sex-Specific Neuron Types and the Ascending Projection Neuron Repertoire

**DOI:** 10.64898/2026.06.11.731752

**Authors:** Liliana Cano-Gomez, Helen Poldsam, Satoshi Ishishita, Ethan Irrobali, Helen C. Lai, Allan-Hermann Pool

## Abstract

The spinal cord gives rise to central somatosensation and orchestrates autonomic and motor control. How its cellular diversity achieves functions across sex, longitudinal axis and links the spinal cord to the brain remains poorly understood. Here we used retrograde viral tracing with spatial transcriptomics and multiomic profiling of over 750,000 mouse spinal cord neurons to define its output logic and functional cellular repertoire. We identified 78 ascending spinal projection neuron classes that link it to hind-, mid-, and forebrain centers for itch, touch, and pain revealing its output wiring logic. Furthermore, we characterized >500 anatomically and transcriptomically distinct neuron types with extensive (∼55%) rostro-caudal specialization as well as first sex specific spinal neurons. Finally, we assembled a cell-type-to-phenotype map of spinal output neurons based on genetically defined interventions. Consequently, we deliver an anatomically and functionally annotated atlas of the adult mouse spinal cord to guide future interrogation of the organ.

## Introduction

The spinal cord (SC) is a sensory-motor structure that gives rise to central somatosensation through its ascending projections as well as orchestrates motor and autonomic functions. These roles are achieved by genetically defined and functionally specialized neuron types. How cellular diversity in the spinal cord implements these functions however remains intensely studied yet poorly understood. The advent of single-cell gene expression profiling has established a transcriptomic cell-type based framework for characterizing the cellular building blocks of SC circuits. Multiple groups have profiled mouse, primate and human spinal cords with single-cell^1–3^ or single-nucleus RNA-sequencing^4–9^ and more recently with spatial transcriptomic methods^9–12^. Surprisingly, these efforts have yielded widely varying estimates of neuronal diversity ranging from 17 to 50 transcriptomic neuron types in the adult mammalian spinal cord. These figures sit uneasily with focused studies of restricted populations such as cholinergic neurons which constitute <2% of spinal neurons yet resolve >30 transcriptomic types^13,14^. Therefore, the granularity at which SC neurons should be analyzed to capture functional logic of sensory-motor processing and how distinct functions map to the underlying neuronal diversity remains largely unclear. A central challenge therefore is to understand how molecularly defined spinal cord neuron types are anatomically organized, and how this organization supports region-specific roles, sexually dimorphic functions, and gives rise to ascending output pathways.

Spinal cord function changes markedly along the rostro-caudal axis as organ systems, muscle groups and autonomic targets alter along this dimension. How SC cellular diversity meets these varying demands is incompletely understood but has recently been explored for the cholinergic system. These studies demonstrated that mammals have evolved new transcriptomic neuron types to implement rostro-caudally specific functions such as over a dozen specialized visceral motor neuron classes^13,14^. These cells control internal organ systems and are mostly confined to the thoracic and sacral spinal cord. How the glutamatergic and GABAergic systems are adjusted remains much less clear. One recent study from humans suggests that neuron types are largely proportionally represented across the spinal cord regions^9^ whereas another rodent study showed dramatic rostro-caudal differences in almost all neuron types^12^. The neural substrates underlying region-specific spinal functions therefore remain incompletely defined.

The spinal cord also implements sexually dimorphic functions, including reproductive reflexes, behaviors and sex-biased pain processing^15–19^. Sex-specific neuron types are well characterized in higher brain centers controlling social and reproductive behaviors^20^, yet despite multiple atlas-scale efforts, no transcriptomically distinct sex-specific neuron types have been reported in the mammalian spinal cord^9,13,21^. Existing studies have instead identified dimorphic gene expression within shared neuron classes, modest abundance biases, or sex-dependent injury responses in glia^12,19,21,22^, leaving open whether sex-dependent spinal functions are encoded by transcriptomically distinct neurons or by graded modulation of common ones.

Most central somatosensation arises from ascending spinal projection neurons (SPNs) that relay nociceptive, thermal, mechanical, itch, and proprioceptive signals to brainstem, midbrain, and forebrain targets and drive supraspinal behaviors^23–25^. The diversity and molecular properties differentiating SPNs from the rest of SC cell-types remain intensely studied. Work from many labs has identified genetic markers that label SPNs implicated in mechanical itch (Calcrl)^26^, chemical itch (Tacr1, Grpr, Cck)^26–28^, heat (Tacr1, Lypd1, Tac1, Gpr83, Nptx2, Crh, Cck, Phox2a)^1,27,29–33^, mechanical (Tacr1, Tac1, Gpr83, Phox2a)^30–32,34^ and visceral pain (Brn3a)^35^, warmth (Tacr1, Gpr83)^33,34^ and cold (Phox2a, Tacr1)^31,33^ sensations as well as proprioception (Gdnf, Slc17a7)^36–39^ and touch (Zic2)^40^. However, whether individual markers label a single transcriptomic neuron class or broader populations and how the markers relate to each other is largely unclear. Surprisingly, published SC transcriptomic atlases have mostly failed to disambiguate this issue. To our knowledge, only four studies have attempted to identify the transcriptomic identity of projection neurons in restricted spinal cord domains or developmental lineages finding between one to twelve candidate excitatory transcriptomic spinal projection neuron classes^1,31,38,41^. In contrast, a recent comprehensive anatomical analysis of spinal projection neurons in the cervical spinal cord revealed >20 structural classes based on central projection targets^42^ suggesting considerably higher SPN diversity. This has led to speculations that adult spinal projection neurons are not molecularly distinct from other SC cell types^43^ or that they may be absent in existing atlases due to tissue dissociation artifacts^44^. As a result, the full SPN repertoire, somatosensory coding logic and their unique molecular features remain surprisingly poorly understood.

To address these outstanding issues, we used deep single-cell multiomic profiling and single-molecule spatial transcriptomics with retrograde viral tracing to characterize the full rostro-caudal and sex-specific cellular diversity as well as the cellular output logic of the spinal cord. We identified the cellular substrates that link the spinal cord to the central brain. Furthermore, we characterized the first sex specific neuron types in the spinal cord and generated a cell-type-to-phenotype map of the spinal cord output neurons. Finally, we devised an interactive online atlas for exploring molecular identity, anatomic properties and transcriptomic projection neuron repertoire of the full adult mouse spinal cord. As a result, we deliver a comprehensive transcriptomically, anatomically and projectionally annotated cell-type framework to guide functional dissection and annotation of the spinal cord as well as illuminate cellular underpinnings of spinal cord disease.

## Results

### A multimodal atlas resolves >500 neuron types and regionalization in the adult spinal cord

To define the cellular composition of the adult mouse spinal cord at high resolution, we combined two complementary profiling modalities. We first dissected the four major rostro-caudal regions (cervical, thoracic, lumbar, and sacral) from male and female mice (n=3, per region per sex), isolated NeuN+ nuclei by flow cytometry, and profiled gene expression and chromatin accessibility in parallel using the 10x Genomics Multiome platform. The resulting dataset comprised 169,345 high-quality neuronal nuclei (**Figure 1A**). Crude Leiden clustering segregated these neurons into 57 broad marker domains—28 glutamatergic, 24 GABAergic, and 5 cholinergic—each defined by a conserved broad marker gene (**Figure 1B-D**). We treat these "groups" as an intermediate hierarchical level corresponding to the family or group ranks in published taxonomies^7,11^. For clarity, we named them based on their neurotransmitter profile and an evolutionarily conserved broad marker consolidated with recent pan-mammalian and human SC atlases^7,45,46^ (**Figure S1A**). Iterative unsupervised clustering within each group resolved 2–35 neuron types per group, yielding 516 transcriptomic neuron types across the full rostro-caudal axis (**Figure 1B-E**). Each type is named by a three-part scheme combining neurotransmitter (GLUT/GABA/CHOL), broad group marker, and finally a narrow marker gene which uniquely identified the neuron type within its group/broad marker domain (**Table S1**). To facilitate the exploration of the molecular, anatomical and projectome data for the spinal cord neuronal diversity, we generated an interactive online platform (https://spinal-cord-explorer.shinyapps.io/mouse/) summarizing the results of our cross-modal SC analysis.

**Figure 1:**
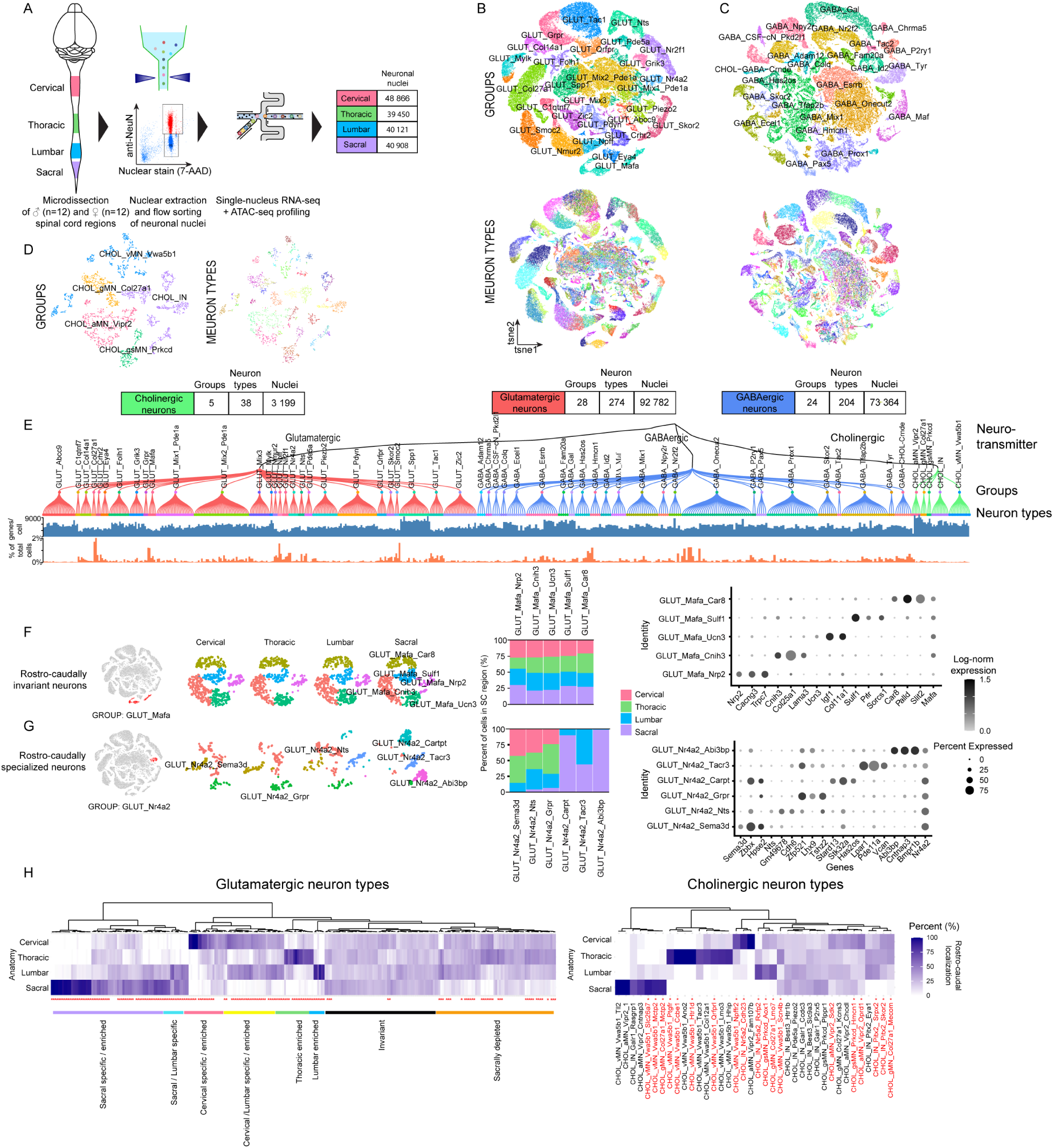
Multiomic characterization of rostro-caudal neuronal diversity in the adult mouse spinal cord. (A). Multiomic (scRNA-seq/sn-ATAC-seq) profiling of neural diversity in the adult mouse spinal cord. (B) tSNE embedding of glutamatergic neurons (n= 92 782 cells) at the group (up) and neuron type level (down). (C) tSNE embedding of GABAergic neurons (n= 73 364 cells) at the group (up) and neuron type level (down). (D) tSNE embedding of all cholinergic neurons (n= 3199 cells) at the group (left) and neuron type level (right). (E) Transcriptomic taxonomy dendrogram of 516 spinal cord neuron types organized into 57 transcriptomic groups and three neurotransmitter classes. (F) Rostro-caudally invariant neuron types in the GLUT_Mafa group; tSNE embedding of GLUT_Mafa neuron types in SC rostro-caudal regions (left, n=2826 cells); rostro-caudal distribution of GLUT_Mafa neuron types from regionally balanced sample (middle, n= 2608 cells); dotplot of neuron type specific markers for GLUT_Mafa constituent neuron types (right). (G) Rostro-caudally specialized neuron types in the GLUT_Nr4a2 group; tSNE embedding of GLUT_Nr4a2 neuron types in SC rostro-caudal regions (left, n=1110 cells); rostro-caudal distribution of GLUT_Nr4a2 neuron types from regionally balanced sample (middle, n=920 cells); dotplot of neuron type specific markers for GLUT_Nr4a2 constituent cell types (right). (H) Rostro-caudal distribution of glutamatergic (left) and cholinergic (right) neuron types in the mouse spinal cord. Neuron types marked by red asterisk/text are rostro-caudally significantly specialized (three-fold enrichment between at least two rostro-caudal regions, adjusted p<0.05 DeSeq2 and adjusted p<0.05 with beta regression, see **Table S1** for glutamatergic neuron identity labels).

We next asked whether group-level analysis is sufficient to capture the known regional and sex-specific specialization of the spinal cord. Transcriptomic groups capture functionally meaningful cell assemblies with sometimes known underlying heterogeneity. For example, visceral motor neurons (CHOL_vMN_Vwa5b1) fall into sympathetic and parasympathetic preganglionic visceral motor neuron classes with diverse independent functions regulating autonomic control of internal organs^13,14^. Each transcriptomic group was defined by characteristic marker gene expression and group-specific open-chromatin profiles (**Figure S1B–E**). With regionally balanced sampling, group-level abundances were uniform across rostro-caudal segments for all but one glutamatergic and GABAergic groups. The only robust regional biases recovered the well-known cervico-lumbar enrichment of skeletal motor neurons (CHOL_aMN_Vipr2, CHOL_gMN_Col27a1, CHOL_gsMN_Prkcd) and thoraco-sacral enrichment of visceral motor neurons (CHOL_vMN_Vwa5b1) (≥3-fold difference between any two regions, adjusted p<0.05, DESeq2 and beta-regression; Figure S2A–C) (**Figure S2A-C**). Group-level analysis on sex-balanced samples detected no sexual dimorphism whatsoever (**Figure S2D-F**). These results indicate that conventional group-resolution clustering masks much of the cellular specialization in the spinal cord.

In contrast, neuron type–level analysis revealed striking rostro-caudal organization. We identified two contrasting patterns of rostro-caudal regionalization. On one extreme, we observed neuron types that were invariant and present in equal proportions at all rostro-caudal levels such as neuron types belonging to the GLUT_Mafa group (**Figure 1F**). On the other extreme, we observed strongly regionally biased neuron types such as neuron classes belonging to the GLUT_Nr4a2 group (**Figure 1G**). Extending this analysis to all neuron types identified 284 high confidence rostro-caudally specialized neuron types (3-fold difference between any two regions, adjusted p<0.05 for DeSeq2 & beta-regression). Collectively these data suggest that slightly over half (55%) of all neuron types in the spinal cord are regionally specific (**Figure 1G**, **S2**) revealing candidate neural substrates for rostro-caudally varying functions beyond previously known cholinergic neuron types.

### Spatial transcriptomics anatomically annotates spinal cellular diversity

To validate this neuronal taxonomy with an independent modality and to place each neuron type in anatomical context, we performed single-molecule spatial transcriptomics on the 10x Genomics Xenium platform. We designed a 480 gene panel including key molecular markers for major cell classes, neuronal groups and types as well as ectopic sequences for retrograde tracing (**Table S2**). Profiling 523 spinal cord sections from cervical, thoracic, lumbar, and sacral regions of 7 male and 7 female mice yielded 616,599 neurons and 2,081,760 non-neuronal cells (**Figure 2A**). Unsupervised clustering identified 18 molecularly and anatomically distinct major cell classes (**Figure 2B-D**) with most showing non-uniform distribution across the cord. Several related cell classes showed segregation between white and gray matter regions including unexpectedly diverse vasculature associated cell classes (venous endocytes, arterial endocytes T1 and T2, VSMCs, pericytes), white matter oligodendrocytes (WM) and non-selective oligodendrocytes (NS) as well as the previously documented WM Astrocytes and GM Astrocytes. Several detected cell-classes were associated with the spinal cord through input/output nerves (myelinating and non-myelinating Remak Schwann cells), circulation (macrophages) or meninges (pial and arachnoid fibroblasts) (**Figure 2D**).

**Figure 2:**
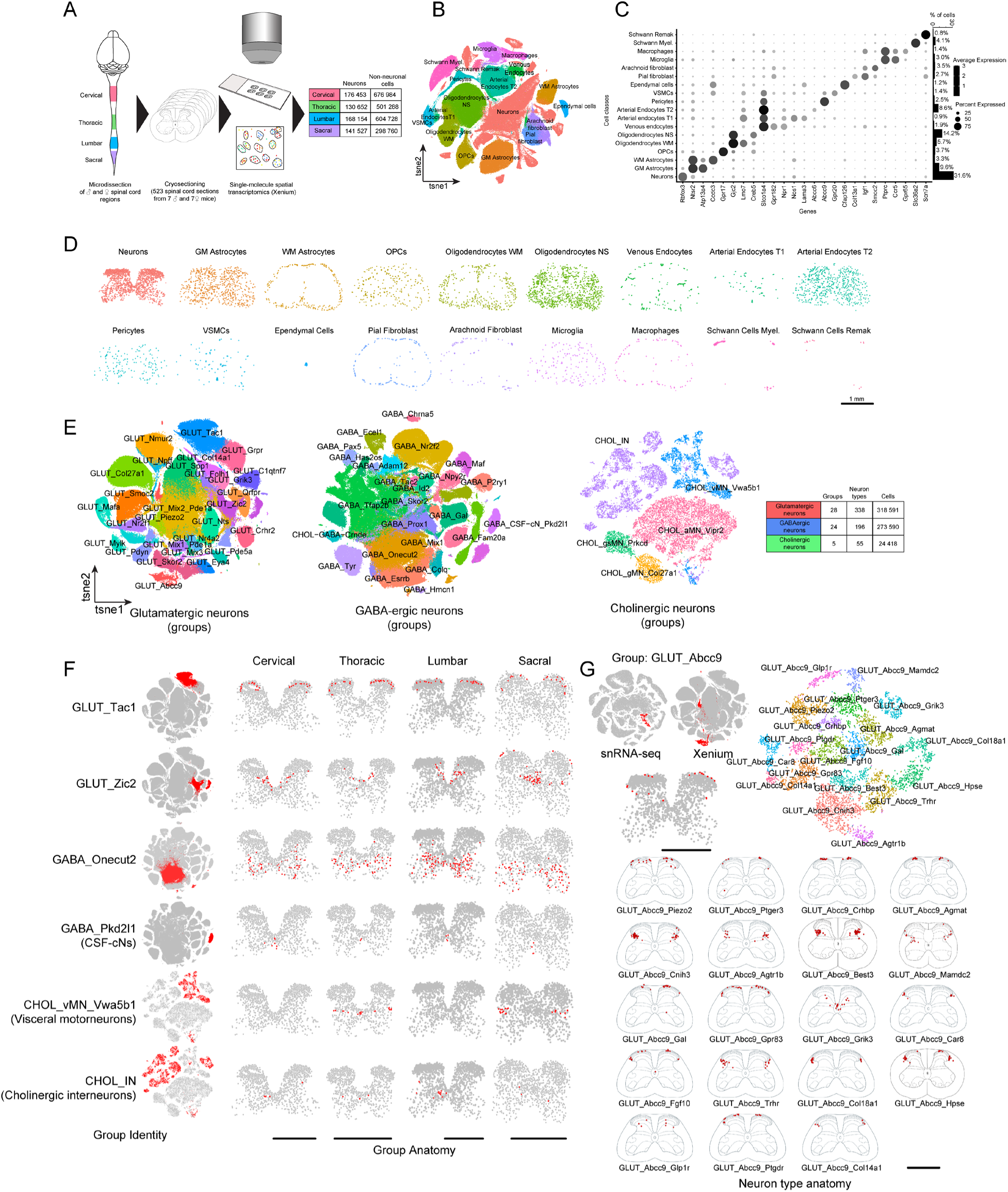
Anatomical annotation of major cell classes, neuronal groups and types in the mouse spinal cord. (A) Generation of single-molecule spatial transcriptomic data from rostro-caudal regions of the adult mouse spinal cord. (B) Major cell classes of the adult mouse spinal cord (tSNE embedding of 383 308 cells from 68 mouse spinal cords). (C) Dotplot of cell class markers (left, log_2_-transformed expression) and bar plot of cell class prevalences (right). (D) Anatomical distribution of major cell classes in a representative 10 µm lumbar spinal cord section (scale bar = 1mm). (E). tSNE embedding of 318 591 glutamatergic (left), 273 590 GABAergic (middle) and 24 418 cholinergic (right) neurons profiled by 10x Genomics Xenium platform (color coded group level identities). (F) Anatomical annotation of transcriptomic groups; group highlighted in tSNE embedding (left, red), highlighted group neurons (red, right) in representative 10-µm adult mouse spinal cord sections at different rostro-caudal levels (scale bar = 1mm). (G) Anatomical annotation of GLUT_Abcc9 within-group neuron types in the mouse spinal cord. tSNE embedding of glutamatergic neurons profiled by multiomic (up left) and spatial transcriptomic (up middle) profiling with GLUT_Abcc9 neurons highlighted in red. GLUT_Abcc9 group neurons in lumbar spinal cord (highlight in red, 10-µm section, up second row). tSNE embedding of GLUT_Abcc9 neurons with color-coded neuron types (up right, n= 7542 cells). Anatomical atlas registered (L5 lumbar spinal cord or non-lumbar reference if absent in lumbar cord) and aggregated (from 10 male and 10 female spinal cord sections) neuron type locations (lower panels, scale bar = 1 mm).

We aligned the spatial and multiomic taxonomies through canonical correlation analysis and label transfer of group identities (**Figure 2E**, **S3A**). The spatial data reproduced the absence of sexual dimorphism and the largely uniform rostro-caudal distribution of glutamatergic and GABAergic groups, while again recovering motor neuron region biases (**Figure S3B–G**). This confirms that more granular neuron type level analysis is necessary to capture most cellular mechanisms underlying sexually dimorphic and other rostro-caudally varying functions. Iterative subclustering within spatial groups resolved 339 glutamatergic, 196 GABAergic, and 56 cholinergic neuron types, with selective markers for each (**Table S3**, https://spinal-cord-explorer.shinyapps.io/mouse/). Importantly, neuron type identities were largely concordant across multiomic and spatial modalities, although a subset were uniquely resolved in one modality—reflecting the higher genetic dimensionality of multiomics, the lower dissociation-induced cell loss of in situ profiling, and differences in anatomical sampling (**Methods**).

Spatial location data enabled systematic anatomical annotation of all 500+ neuron types. Each transcriptomic group occupied a stereotyped territory ranging from single-laminar (e.g., GLUT_Grpr in lamina I; GLUT_Tac1) to broadly dispersed distributions (e.g., GLUT_Abcc9, GABA_Esrrb; **Figure 2F**, **S4**). As anatomical location is essential for interpreting spinal neuron identity, we further developed a spatial atlas registration pipeline that we used to aggregate spatial distribution of individual neuron types across several sections thus allowing robust anatomical characterization of neuron type level anatomy (https://spinal-cord-explorer.shinyapps.io/mouse/). Strikingly, anatomically dispersed groups such as GLUT_Abcc9 comprised neuron types that subdivide the group’s footprint into spatially tiling subdomains, revealing previously unappreciated genetic-anatomical structure within transcriptomic groups (Figure 2G). The same neuron type–level analysis recovered the full diversity of cholinergic neurons, including previously characterized motor neuron classes and additional cholinergic interneuron and visceral motor neuron types (**Figure S5**). Collectively, our analysis reveals several fold higher diversity among the spinal cord neuron types demonstrating over 500 genetically and anatomically distinct neuron classes across the full rostro-caudal axis. Importantly we demonstrate that many transcriptomic entities that were characterized as individual neuron types in previous atlases comprise anatomically and genetically highly divergent neuron types. These data argue for a more granular analysis of spinal cord circuits and underscore the importance of incorporating both transcriptomic and anatomic data to resolve cellular diversity in neuronal tissue.

### Sex specific neuron types in the spinal cord

Because group-level analysis detected no sexual dimorphism, we asked whether sex-specific cellular identity emerges at neuron type resolution. Sex-balanced sampling of the spatial transcriptomic data, followed by rank-ordered analysis of male/female bias, identified 4 male-specific and 3 female-specific neuron types that were significantly sex-biased (n=7 male and 7 female mice; DeSeq2, adjusted p<0.05; **Figure 3A,B**). Interestingly, sex specificity was restricted to cholinergic and glutamatergic neurons with no significant sex biases among GABAergic neurons (**Figure S6A-C**). Moreover, cholinergic sex specific neuron types belonged to the visceral motor neuron group (CHOL_vMN_Vwa5b1) and were defined by non-sex chromosome derived marker gene expression (**Figure 3C,D**). Specifically, the male specific CHOL_vMN_Vwa5b1_Tacr3 neurons were located in the lower thoracic spinal cord. Furthermore, we observed a male specific (CHOL_vMN_Vwa5b1_Grp) and a female specific (CHOL_vMN_Vwa5b1_Col14a1) visceral motor neuron type in the lateral horn of the sacral spinal cord (**Figure 3E**). Intriguingly, thoracic and sacral autonomic systems have been suggested to control emission of gametes and erectile/secretory functions, respectively^47^ placing these neurons as candidate substrates for sexually dimorphic autonomic control of reproduction.

**Figure 3:**
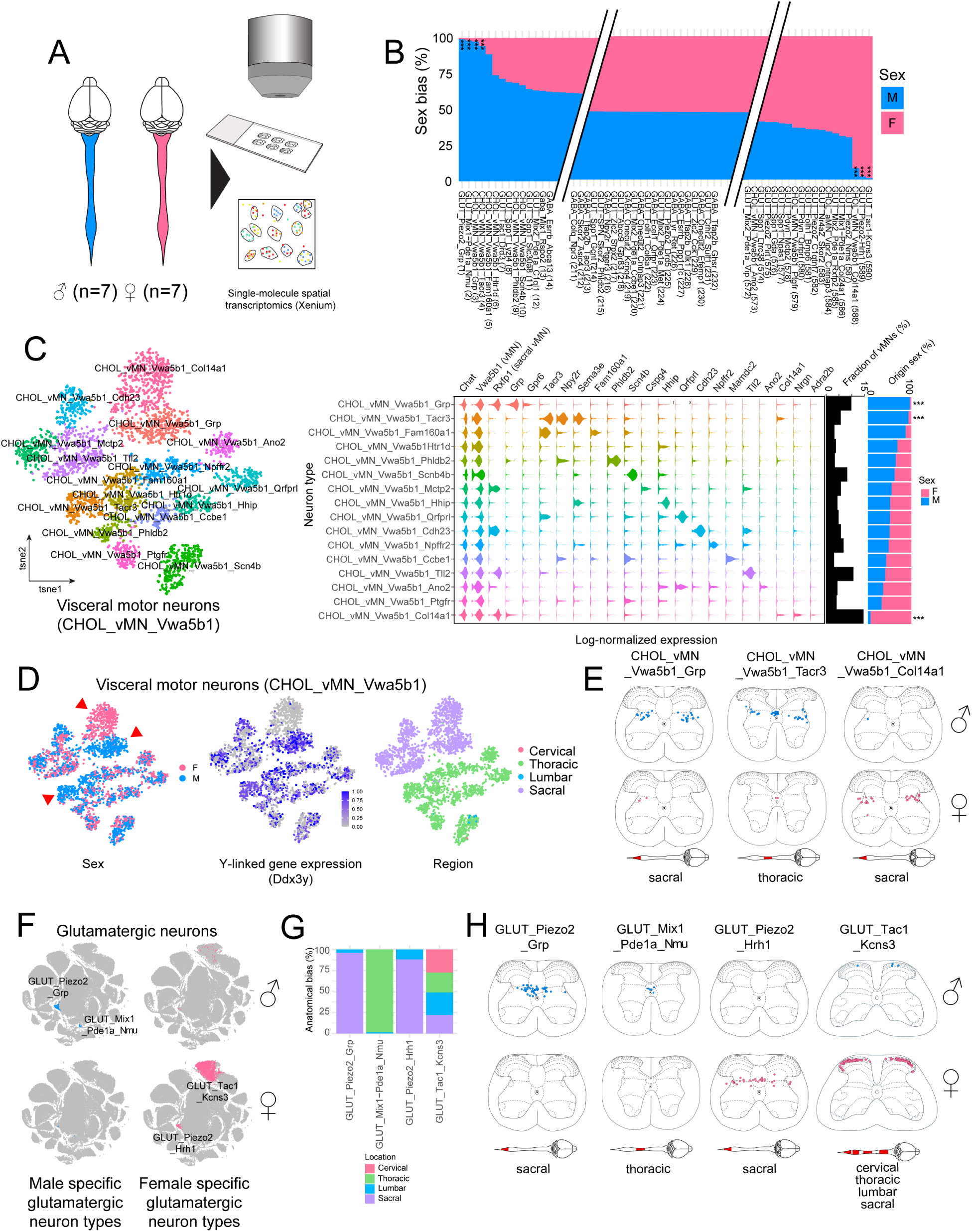
Sex specific neuron types in the spinal cord. (A) Spatial transcriptomic profiling of sex specific differences in the adult mouse spinal cord. (B) Rank-ordered neuron types based on male/female bias from sex balanced sample (sex bias estimated with DeSeq2, comparing models including or excluding sex, n=7 mice per sex, n= 303 835 cells from each sex, *** adjusted p<0.001). (C) tSNE embedding of visceral motor neuron types (group CHOL_vMN_V2a5b1, left, n= 3975 cells) and violin plot of cell-type specific marker genes (right, log_e_ transformed expression, sex bias estimated with DeSeq2, comparing models including or excluding sex, *** adjusted p<0.001). (D) tSNE embedding of visceral motor neurons (CHOL_vMN_V2a5b1) with sex identity (left), expression of Y-linked male specific Ddx3y gene (middle) and color-coded rostro-caudal location of cells (right). Red triangles indicate sexually dimorphic cholinergic neuron types. (E) Anatomical location of sex-specific visceral motor-neuron types aggregated from 10 male (up) and 10 female (below) spinal cord sections. (F) tSNE embedding of glutamatergic neurons (n= 318 778 cells), with highlighted male specific (blue) and female specific (pink) neuron types. (G) Rostro-caudal distribution of sex-specific glutamatergic neurons. (H) Anatomical location of sex-specific visceral motor-neuron types aggregated from 10 male (up) and 10 female (below) spinal cord sections.

We also observed four types of sexually dimorphic glutamatergic neurons. Three of these (GLUT_Piezo2_Grp, GLUT_Piezo2_Hrh1, GLUT_Mix1_Pde1a_Nmu) were restricted to either sacral or thoracic spinal cord and one (GLUT_Tac1_Kcns3) showing a wider distribution (**Figure 3F-H**). Importantly, most of these sex biased cell types were evident in both multiomic as well as Xenium datasets (**Figure S6D-G**). Moreover, we observed some neuron types that displayed moderate sex biases but which were statistically insignificant with the vast majority of neuron types showing no biased sex distribution (**Figure 3B**). In sum, these experiments identify to our knowledge the first sex specific transcriptomic neuron types in the mammalian spinal cord that constitute candidate neural substrates for sexually dimorphic behaviors.

### Identification of Ascending Spinal Projection Neuron Repertoire

The spinal cord gives rise to central somatosensory perception through ascending spinal projection neurons that relay information about pain, itch, touch, temperature, and proprioception to higher order brain centers. The transcriptomic identity, wiring logic and functional role of most SPNs however remain poorly understood. Here, we used multiplexed cell-type resolved retrograde viral tracing with spatial transcriptomics to identify the ascending projection neuron repertoire throughout the adult spinal cord (**Figure 4A**). To this end, we injected retrograde AAV tracers to major spinal cord output structures in the forebrain (mediodorsal thalamus - MD, ventral posterolateral nucleus of the thalamus - VPL, posterior triangular thalamic nucleus - PoT), midbrain (periaqueductal gray - PAG, superior colliculus - SCOL) and hindbrain (lateral parabrachial nucleus - LPBN, cerebellum - CEB, caudal ventrolateral medulla - CVLM and dorsal column nuclei - DCN) in both male and female mice (n=2 male and 2 female mice per tracing target). Retrograde tracing led to robust labeling of projection neurons (**Figure 4B**, **S7**) that were enriched in 13 out of 28 glutamatergic groups (**Figure 4C**). De-aggregating the data at the neuron type level revealed that 78 out of 338 glutamatergic neuron types link the spinal cord to one or more central brain targets (**Figure 4D**, **S8**, permutation test with Bonferroni adjusted p<0.05) thus defining the transcriptomic identity of the mouse spinal cord output neurons. Critically, we show that spinal cord output neurons are transcriptomically distinct neuron types clearly discernible from the rest of spinal neural diversity.

**Figure 4:**
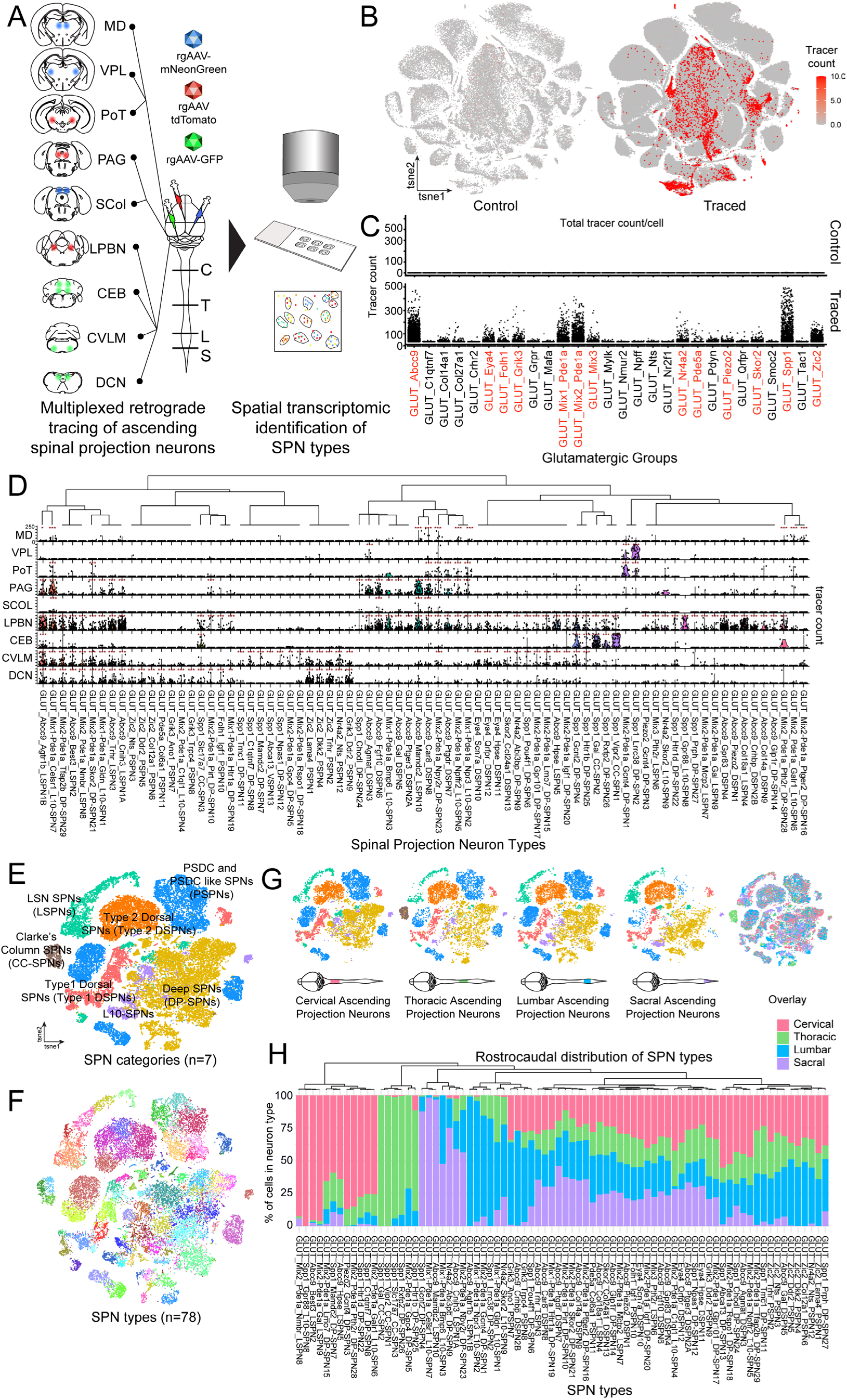
Retrograde viral tracing from major spinal cord output structures identifies the ascending spinal projection neuron repertoire. (A) Multiplexed cell-type resolved retrograde tracing of ascending spinal projection neurons (rgAAV – retrograde AAV). (B) Feature plot of total retrograde tracer transcript count (tdTomato, GFP, mNeonGreen) in glutamatergic neurons in non-traced (left, n=2 mice, 40 281 cells) and traced animals (right, n=12 mice, 278 310 cells). (C) Scatter plot of total tracer transcript count in glutamatergic groups from non-traced and traced animals. Groups containing statistically significantly labeled neuron types highlighted in red. (D) Scatter/violin plot of tracer transcript counts in spinal projection neuron types per tracing target structure (permutation test with Bonferroni adjusted p-values, * p<0.05, ** p<0.01, *** p < 0.001, n=4 animals per traced structure). (E) tSNE embedding of spinal projection neuron types (n=14 animals, 39 527 cells), color coded by anatomical category. (F) tSNE embedding of spinal projection neuron types color coded by neuron type. (G) Rostro-caudal distribution of spinal projection neuron categories. Overlay of rostro-caudal regional origin (color-code same as panel H). (H) Rostro-caudal distribution of spinal projection neuron types.

To facilitate the analysis of ascending SPNs, we extracted the projection neuron types from the rest of the atlas and performed detailed molecular and anatomic characterization of the system. Crude unsupervised clustering of these neurons identified 7 anatomically and molecularly distinct categories of SPNs that shared within-category anatomical location and partially molecular features (**Figure 4E**). These included two categories in the superficial dorsal horn of the spinal cord (Type 1 and Type 2 DSPNs), one category aggregating lateral spinal nucleus (LSN) spinal projection neurons (LSPNs) and a category of Clarke’s Column SPNs (CC-SPNs) projecting to the cerebellum. Furthermore, we observed a molecularly diverse category of dorsal column projecting SPNs that resemble touch information relaying post-synaptic dorsal column projection neurons^48,49^ (PSPNs), peri-central canal L10-SPNs and finally a heterogenous group of deep SPNs (DP-SPNs). Critically, all the categories comprise diverse neuron types totaling 78 SPN classes (**Figure 4F**). Only Clarke’s column and PSPN categories showed strong rostro-caudal regionalization with neurons only in the thoracic segment and depleted in the sacral regions, respectively. However, individual SPN types within all categories showed diverse anatomical regionalization along the rostro-caudal axis with ∼2/5 of the SPN types present throughout the spinal cord. The rest localized selectively in 1-3 regions (**Figure 4H**). Collectively, these data demonstrate several fold higher molecular and anatomic diversity of ascending spinal projection neurons and that different rostro-caudal SC regions are linked to the central brain in part by transcriptomically distinct neuron types.

Based on the above experiments, we compiled a molecular-anatomic catalog of spinal cord SPNs (**Figure 5**, **S9**). Importantly, we find that although many ascending neuron types share anatomical features, the underlying cells are molecularly distinct, even within the constrained 480 spatial gene set. Moreover, these data demonstrate the transcriptomic neuron type composition of previously identified SPN markers and how these cell populations relate to each other. Specifically, we found that the Tac1+ SPNs implicated in pain coping behaviors are cellularly highly diverse labeling the majority of ascending neuron types in the superficial dorsal horn and LSN. We also found that adult Phox2a and Gpr83+ SPN populations implicated in pain processing were non-overlapping, while Tacr1 was co-expressed in both based on our snRNA-seq as well as spatial transcriptomic datasets addressing previously contradictory findings^34,41,50^ (**Figure 5**). Furthermore, our tracing experiments reproduce almost all previously known spinal ascending neuron types in expected anatomical locations. Specifically, we identified thoracic Gdnf+/Slc17a7+ ascending Clarke’s column neurons implicated in proprioception with previously unknown anatomical and molecular subtypes (**Figure 5G**). We also found transcriptomic SPN types that directly correspond to the recently characterized Phox2a+ projection neuron classes^44^. For example, GLUT_Abcc9_Agmat_DSPN3 directly corresponds to the putative cold activated ALS3, and GLUT_Abcc9_Ptger3_DSPN2A /GLUT_Abcc9_Crhbp_DSPN2B correspond to the heat activated ALS2 population from the Phox2a+ developmental lineage (**Figure S10B1-5**). Moreover, we found most of the cervical projection neuron types characterized in a recent study^41^ with some differences in anatomic annotation (**Figure S10C1-5**). Importantly, we identified several dozen SPN types outside the Phox2a lineage and previously reported cervical populations including comprehensive characterization of sacral and thoracic SPNs and previously unknown superficial dorsal horn and LSN populations (**Figure 5**, **S9**). Collectively, these data establish a clear molecular/anatomical framework for functional dissection of the ascending SC output projection neuron system.

**Figure 5:**
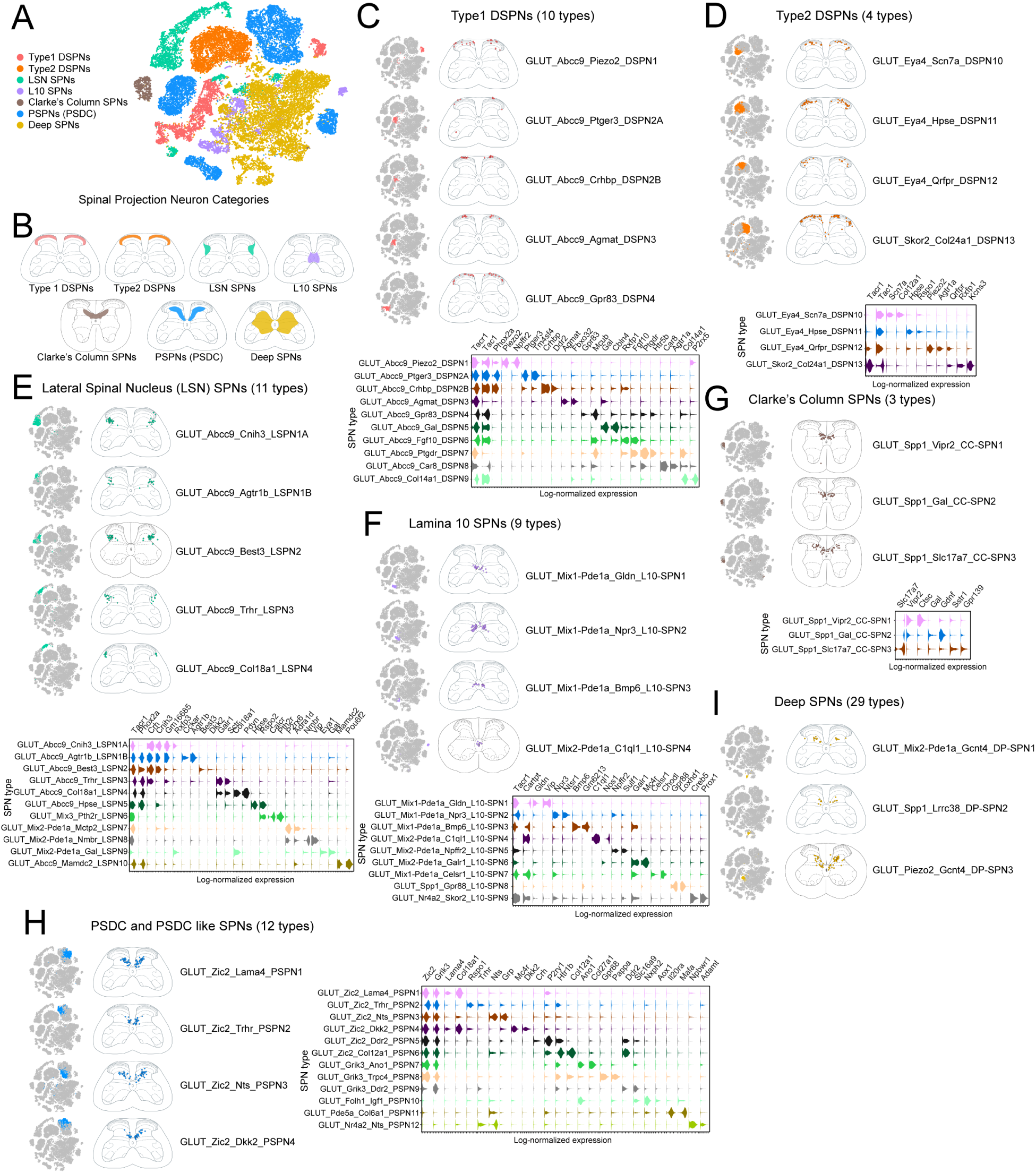
Molecular and anatomic catalog of ascending spinal projection neuron types. (A) tSNE embedding of spinal projection neurons by anatomical category. (B) Anatomical distribution of spinal projection neuron categories in the lumbar spinal cord. (C) Anatomy (up) and molecular markers (violin plot, down) for Type 1 Dorsal horn Spinal Projection Neurons (Type 1 DSPNs, DSPN1-9). Continued in Figure S9A. (D). Anatomy (up) and molecular markers (violin plot, down) for Type 2 Dorsal horn Spinal Projection Neurons (Type2 DSPNs, DSPN10-13). (E) Anatomy (up) and molecular markers (violin plot, down) for LSN SPNs (LSPNs). Continued in Figure S9B. (F) Anatomy (up) and molecular markers (violin plot, down) for Lamina 10 SPNs (L10-SPNs). Continued in Figure S9C. (G) Anatomy (up) and molecular markers (violin plot, down) for Clarke’s Column SPNs (CC-SPNs). (H) Anatomy (up) and molecular markers (violin plot, down) for postsynaptic dorsal column (PSDC) and PSDC-like SPNs (PSPNs). Continued in Figure S9D. (I) Anatomy (up) and molecular markers (violin plot, down) for Deep SPNs (DP-SPNs). Continued in Figure S9E. For (C)-(H) cell type anatomy registered from 20 spinal cord 10 µm sections to reference lumbar (L5) spinal cord atlas or other (cervical, thoracic, sacral) if cell type was not present in lumbar spinal cord.

### Wiring logic of the ascending spinal output

Spinal projection neurons innervate dozens of supraspinal targets, with axon collateralization patterns ranging from highly restricted to broadly distributed^24,42,51^. How these anatomical projection patterns relate to transcriptomic identity of projection neurons remains poorly understood. To relate molecular identity to brain-wide wiring, we used the multiplexed retrograde tracing data to construct an SPN connectivity matrix (**Figure 6A**) identifying both highly restricted as well as widely collateralizing SPN classes recapitulating the wiring diversity of an earlier anatomical survey^42^. Specifically, we observed many SPN subtypes with projection targets restricted to the brainstem, medial or lateral thalamic structures (**Figure 6A**). In contrast, we also observed dozens of SPN classes with broad arborization from brainstem to thalamus.

**Figure 6:**
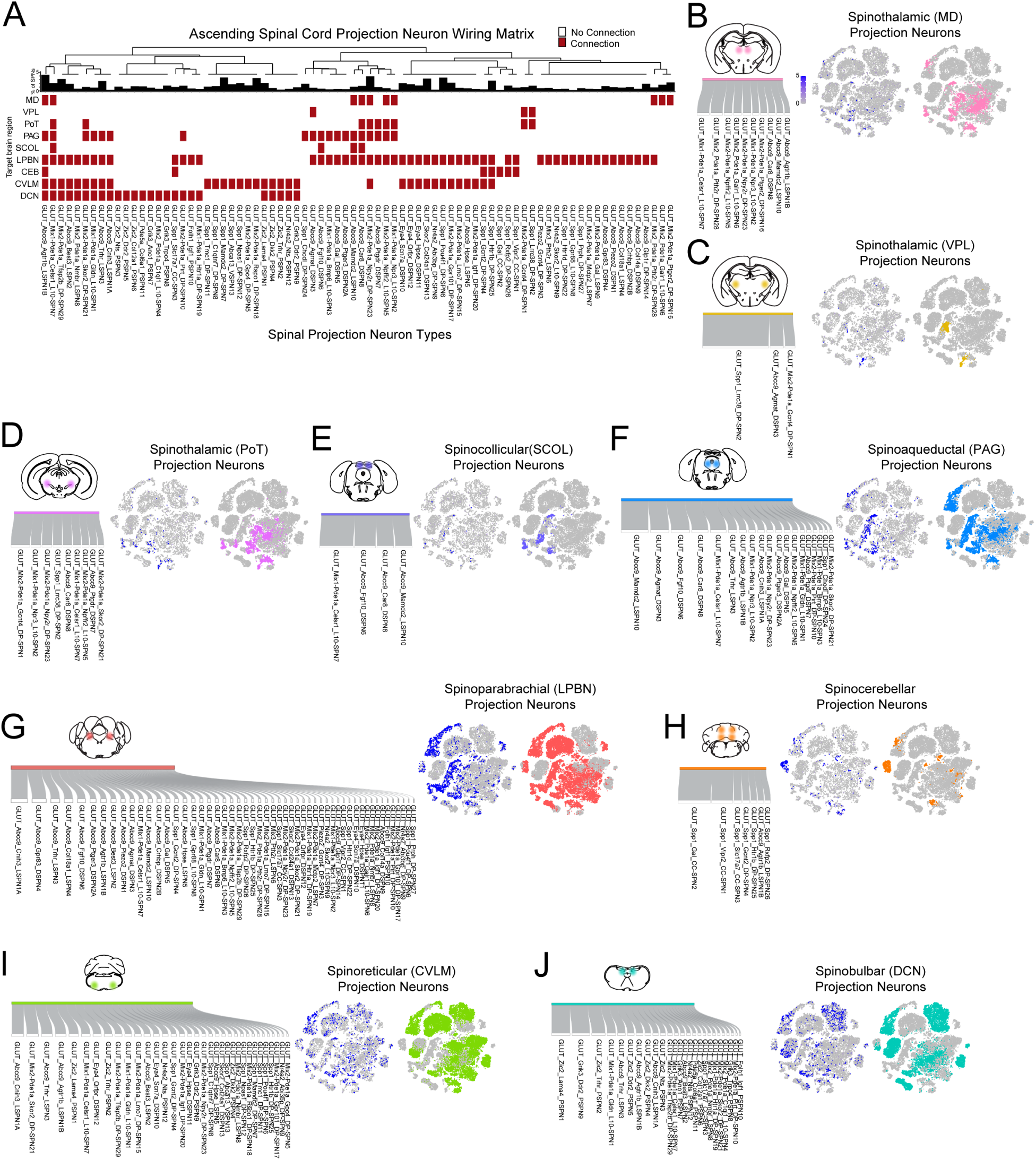
Wiring logic of the ascending spinal projectome. (A) Spinal cord connectivity matrix for identified spinal projection neuron types based on connectivity analysis in Figure 4. Bar graph (up) showing proportional abundance of SPN types as fraction of all SPNs. Hierarchical clustering dendrogram (up) of SPN class connectivity. (B) – (J). Sankey plot of structure (MD, VPL, PoT, SCOL, PAG, LPBN, CER, CVLM, DCN) projecting SPNs with flow width proportional to labeled SPN cells (left) for given target; Feature plot of tracer (mNeonGreen, tdTomato or GFP) transcript count; structure projecting SPN types (right, highlight). Cell counts: n=16 845 cells for MD, LPBN, DCN; n=9546 for VPL, PAG, CVLM; n=8485 for PoT, SCOL, CEB.

SPN types varied not only in connectivity pattern but also in projection strength to individual targets. To quantify each SPN type’s relative contribution to a given target, we normalized retrogradely labeled cell counts by the total labeled SPN population for each tracing experiment and visualized the resulting proportions as Sankey diagrams (**Figure 6B-J**). For most supraspinal targets, projection strength of individual SPN types varied by more than an order of magnitude, indicating that each downstream structure receives input from a transcriptomically structured ensemble of SPN classes with widely differing weights. Together with the molecular catalog above, these data define the molecular identity, wiring topology, and weighting of the ascending spinal output.

### Cell-type-to-phenotype map of the ascending spinal cord projection neurons

The observed SPN diversity raises the critical question about the functional role of individual SPN types in orchestrating supraspinal somatosensory behaviors. Several studies have used Ca^2+^ imaging, electrophysiological recordings or immediate-early gene based characterization of the somatosensory stimulus tuning of specific genetic marker labeled SPNs^27,28,30,34,35,44,52,53^. Mapping these markers to our SPN atlas revealed the specificity of the SPN markers (**Figure S11A**) as well as the transcriptomic cell-type diversity underlying each SPN marker gene (**Figure S11B**). Many of the SPN markers (e.g. Tacr1, Tac1, Pou4f1, Cck) labeled broad SPN populations as well as were expressed in a number of interneuron classes outside SPNs. This leaves the precise transcriptomic identity of somatosensory stimulus engaged neurons obscure. However, some marker genes such as mechanical pain activated Gpr83+ SPNs^34^, itch activated Grpr+ SPNs^28^, heat activated Phox2a+(high)/Ptger3+/Tm4sf1+^44^ SPNs and putative cold activated Phox2a+(low)/Adamts15+/Gm32828+^44^ neurons label one or a couple of transcriptomic neuron types (**Figure S11B**). This analysis identified discrete transcriptomic neuron classes as candidate neurons driving respective stimulus triggered behaviors based on their somatosensory stimulus tuning. The response properties of most of the uncovered SPN types, however, remain unknown.

Next, we aimed to generate a functional annotation of SPN classes by revealing the SPN combinations that are either necessary or sufficient to orchestrate somatosensory behaviors upon functional intervention. To this end, we included genetic markers in the Xenium gene panel from all the studies that have quantified the effect of SPN restricted genetic, chemo- and/or optogenetic interventions on central pain, itch, touch and temperature behaviors^26,30,31,34,40^ to compile a cell-type-to-phenotype map of SPN types (**Figure 7**). All SPN genetic markers yielding robust to intermediate pain phenotypes upon gain or loss of function manipulations (Tacr1^34^, Tac1^30^, Phox2a^31^) label a third to a majority of Type 1 DSPNs in the lamina I-II superficial dorsal horn and LSN SPNs (LSPNs) implicating these broad SPN categories in pain processing (**Figure 7A**). However, these categories comprise many SPN types and it remains unclear whether single or distinct combinations of SPNs within or across SPN categories need to be engaged to execute pain behaviors. In contrast, Zic2+ SPNs which regulate light touch sensitivity but do not modulate pain responses^40^ label an orthogonal combination of SPNs almost exclusively in the PSPN category. Again, the Zic2 labeled PSPNs are molecularly heterogenous. Similarly, SPN markers yielding cold and itch phenotypes significantly narrow down the transcriptomic SPN types underlying these sensory modalities (**Figure 7B-C**). Finally, we observed a transcriptomic group level segregation between SPNs implicated in pain, itch and temperature sensation as compared to SPNs yielding tactile phenotypes likely reflecting distinct developmental origin of these sensory relay pathways (**Figure 7E 1-5**). Together, the cell-type-to-phenotype map reveals our best existing model of cell-type to phenotype relationships for spinal projection neurons. While still coarse and lacking in fine transcriptomic neuron type resolution, this map implicates DSPN, LSPN, and PSPN categories as principal substrates for distinct supraspinal somatosensory behaviors and, in concert with the molecular-anatomical catalog, establishes a roadmap for neuron type–resolved functional dissection of the ascending spinal outputs.

**Figure 7:**
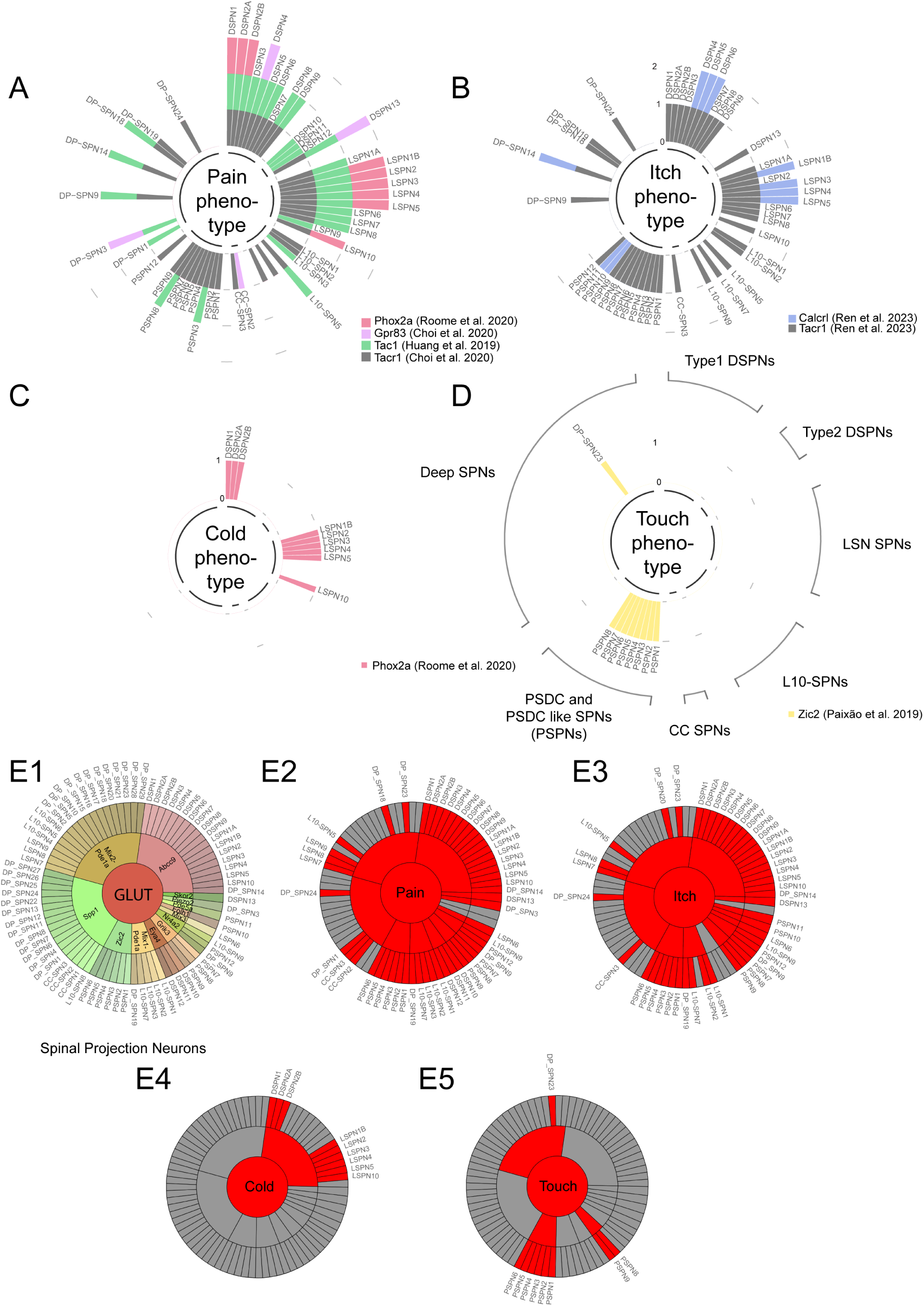
Cell-type-to-phenotype map of the mouse spinal projection neuron system. (A) Expression of genetic markers yielding a pain phenotype in SPN-types and categories upon SPN targeted genetic, chemo- or optogenetic gain and/or loss of function manipulations. Expression is discretized by median thresholding of all positive cells applied to cell-type averaged expression in Xenium profiled SPN atlas. SPN names shortened from Figure 5. SPN types groupd by SPN categories. (B) Genetic markers yielding an itch phenotype upon SPN-targeted manipulation. (C) Genetic markers yielding a cold temperature detection phenotype upon SPN-targeted manipulation. (D) Genetic markers yielding a touch phenotype upon SPN-targeted manipulation. (E1) Dial graph showing group identity of individual SPN types. (E2)-(E5) Highlighted SPN types implicated in pain, itch, cold and touch processing respectively from cell-type-to-phenotype maps in panels (A)-(D).

## Discussion

Cellular diversity is the substrate on which spinal cord function is built, yet the resolution at which spinal circuits should be analyzed has remained an unsettled question. By integrating multiomic single-nucleus profiling, large-scale single-molecule spatial transcriptomics, and multiplexed cell-type-resolved retrograde tracing in both sexes, we provide an anatomy-, transcriptome-, and projectome-annotated atlas of the adult mouse spinal cord. The atlas resolves >500 transcriptomically and anatomically defined neuron types—an order of magnitude beyond prior mammalian atlases ^1,4,7–9,46^. It also reveals previously hidden organizational features of spinal circuits, including the first sex-specific neuron types in the mammalian spinal cord, a 78-class repertoire of ascending projection neurons, and a cell-type-to-phenotype map linking these classes to spinally driven behaviors.

### Resolution matters: granularity is essential to capture spinal cord function

Our results argue strongly that conventional group-level analysis of spinal cord transcriptomes misses much of the biology. At the group level (57 groups), we detected no sexual dimorphism, masked the well-established diversity of visceral motor neurons^13,14^, could not separate spinal projection neurons from interneurons, and obscured projection-neuron diversity by aggregating molecularly and projectionally distinct cells (e.g., within GLUT_Abcc9 and GLUT_Spp1). At neuron type resolution (>500 types), sex-specific cell types emerged, the full cholinergic repertoire was recovered without enrichment, and the molecular identity of ascending output neurons became tractable. Therefore, our results argue that it is essential to analyze spinal neural circuits at the finer neuron type level and demonstrate that with sufficient sampling transcriptomic atlases can sensitively identify functionally important building blocks of spinal circuits. This is also supported by a robustly powered companion study to this paper characterizing spinal cord neural diversity across mammalian species demonstrating over 250 transcriptomic neuron types^45^ as well as recent developmental atlases revealing over a hundred transcriptomic neuron types evident already in early spinal cord development^54^.

### A molecular and wiring framework for the ascending output of the spinal cord

A central finding of our study is that the ascending output of the spinal cord is made up of at least 78 transcriptomically distinct neuron types organized into seven anatomically coherent categories with stereotyped supraspinal wiring. This constitutes a more than five-fold increase from previous efforts to identify transcriptomic identify of SPNs from restricted SC regions^1,41,44^. These spinal cord ascending projection neurons constitute the cellular substrates by which the spinal cord generates central somatosensation and recruits supraspinal circuits during pain, temperature, touch, itch, proprioception, and reproductive behaviors. Because all central pain perception requires intact ascending output, a subset of identified SPNs are also new candidate therapeutic targets for pain^25^.

Several technical reasons explain why this diversity had previously remained elusive in prior atlases. First, we observed a 1.2-3x overrepresentation of the SPN types in our spatial transcriptomic data as compared to multiomic profiling that relies on physical tissue dissociation. This implies that dissociative approaches selectively deplete projection neurons. The second technical limitation obscuring projection neuron identification has been limited cellular sampling which is a well-established problem with detecting rarer neuron types^55,56^ such as SPNs causing them to likely co-cluster with more abundant interneurons in more limited datasets. We overcame this limitation by exhaustive cellular profiling of over 750,000 neurons. Finally, we identified SPNs in the full rostro-caudal span of the spinal cord and found that many SPNs are regionally specific (**Figure 4H**). Critically, we could recognize most of these neuron types across the two profiling modalities (snRNA-seq and Xenium probe based spatial transcriptomics).

The atlas reproduces essentially all SPN populations described in the modern literature. This includes the five lumbar Phox2a+ developmental lineage defined SPN types (ALS1-ALS5) described by Bell and colleagues^44^ (**Figure S10B1-5**). We could clearly distinguish the heat activated ALS2 that corresponds to GLUT_Abcc9_Ptger3_DSPN2A/GLUT_Abcc9_Crhbp_DSPN2B in our data and putative cold activated ALS3 that corresponds to GLUT_Abcc9_Agmat_DSPN3 SPNs (**Figure S10B1-5**). These are all superficial dorsal horn lamina I projection neurons that belong to Type1 DSPN category of SPNs in our nomenclature. This also highlights that SPN types in a single category can be functionally highly heterogenous and need to be independently characterized. Furthermore, we found almost half a dozen other superficial dorsal horn SPN types that do not appear to belong to the Phox2a lineage and whose function is currently unknown with the exception of GLUT_Abcc9_Gpr83_DSPN4 that makes up a significant portion of Gpr83+ projection neurons^34^. Moreover, we identified most of the 12 excitatory cervical spinal projection neurons reported in a recent study by Lin and colleagues^41^ with a few notable differences. We found that the lateral projection neuron cluster (latPN) reported by Lin et al. is in fact a composite cluster of cold and heat sensitive superficial dorsal horn neurons in our (DSPN2A,B and DSPN3) and Bell et al. (ALS2, ALS3) data. This further underscore the importance of careful consideration required in deciding the level of clustering for these datasets and how knowing physiological responses can be indispensable in deciding the appropriate granularity of analysis. Furthermore, Lin et al. dataset lacked lateral spinal nucleus SPNs (LSPN category in our study) where we observed a high diversity (both Phox2a+ and Phox2a-) of SPN types as well as no other L1 superficial dorsal horn SPN types present in our as well as the Phox2a+ lumbar dataset (e.g. ALS1/DSPN1, DSPN5-9). However, Lin et al. data prominently featured SPN types corresponding to our Type 2 DSPNs as well as many of our PSPNs (**Figure S10C1-5**). Interestingly, Lin et al. also reported two dorsal horn populations (dPN2 and dPN3) that were not strongly represented in our data and possibly stem from partially non-overlapping target structures they profiled. Here, future studies with orthogonal methods will prove highly useful to clarify their projection neuron status.

How does SPN diversity give rise to central somatosensory perception and orchestrate behavior? The transcriptomic diversity we characterized suggests that many of the SPN types could serve as functionally specialized labeled lines. Indeed, several *in vivo* calcium imaging studies suggest that Phox2a lineage SPNs show partially non-overlapping heat and cold responses^52,53^ which agrees with immediate early gene analysis and dorsal root ganglion input analysis of candidate cold and heat activated subtypes of Phox2a SPNs^44^. Furthermore, a recent study characterized highly diverse physiological tuning properties of PSDCs^57^ which agrees with the surprising diversity of dorsal column projecting PSPNs in our atlas and could be developmentally defined. However, the physiological response properties and functional role of most SPN types we report remains currently unknown. Due to cellular complexity of the ascending system, new high-throughput methods capable of monitoring activity at transcriptomic neuron type level are required such as immediate early gene based stimulus-to-cell-type mapping approaches^58–60^ or experiments coupling *in vivo* imaging with follow up single-cell transcriptomic profiling^61,62^. Our SPN atlas provides clear molecular-genetic avenues for rational functional dissection of the system.

### Sex differences in the spinal cord

The spinal cord is a major brain center regulating sexually dimorphic behaviors and sensory processing. Despite that, previous spinal cord atlases had surprisingly not reported any sex specific neuron types with only sex biases reported in specific spinal cord regions for a couple of neuron types in a recent spatial transcriptomic survey^12^. This was at odds with earlier work demonstrating both anatomical and neurochemical sex differences in the spinal cord^15,63^. Our finding of seven sex-specific neuron types—predominantly in thoracic and sacral segments—reconciles this gap with the long-standing anatomical and neurochemical evidence for sex differences in the spinal cord. The cholinergic sex-specific types lie within visceral motor neuron pools that regulate lower thoracic and sacral autonomic outflow, providing strong candidates for the cellular control of gamete emission, reproductive tract lubrication, and erectile/secretory function^47^. One of our male specific visceral motor neuron type CHOL_vMN_Vwa5b1_Tacr3 in the lower thoracic spinal cord may potentially correspond to the male enriched visceral motor-neuron population in lumbar spinal cord in a recent study^12^ where subtypes of vMNs were not further resolved. The glutamatergic sex-specific types may represent dedicated relay or modulatory circuits engaged by reproductive sensory inputs, including those from the mammary gland. Somewhat surprisingly, we did not find any sex-specific alpha-motor neurons that have previously been suggested to control erection and locate in the spinal nucleus of the bulbocavernosus^64^. This may be the result of limited sampling and will likely be uncovered in future more dense sampling efforts. Furthermore, we did not identify any sex specific ascending spinal projection neurons which may also require denser sampling. The fact that sex specificity emerges only at neuron type resolution underscores the importance of granularity for studies of dimorphic spinal function.

### Functional annotation of the ascending spinal projection neuron types

Functional annotation of the spinal cord neural diversity remains a formidable challenge and requires a comprehensive transcriptomic cell-type resolved framework to serve as a basis of annotation. Similarly to genes and genomes that benefit from comprehensive phenotypic annotations aggregated by databases such as OMIM^65^ and IMPC^66^, transcriptomic atlases require functional annotation to ascribe human interpretable functions to neuron types. Work from many labs over the past decades has dissected the functional role of genetically defined spinal circuit components in pain^67–69^, itch^70–72^, temperature^73,74^ and touch^40,75,76^ sensation as well as sexual function^63^. Many of these genetic markers define a limited number of transcriptomic neuron types that allow narrowing down and sometimes unambiguous determination of the transcriptomic identity of causal neurons contributing to a specific phenotype. In the current work, we adopted this approach to provide a first pass functional annotation of the spinal cord projection neuron types based on studies deploying SPN targeted functional manipulations (**Figure 7**). The resulting cell-type-to-phenotype map reveals discrete combinations of SPNs that yield significant supraspinal somatosensory phenotypes upon targeted manipulation. Furthermore, analyzing overlap between distinct functional genetic marker domains can start to narrow down the precise transcriptomic identify of causal neurons for individual somatosensory behaviors. This approach should be similarly applicable to interneurons of the spinal cord and thus pave the way for neuron-type-resolved functional understanding of the spinal cord information processing.

### Limitations of the study

Although we aimed to comprehensively sample cellular diversity in the full adult spinal cord, our tissue sampling excluded coccygeal as well as rostral segments of cervical (C1-C2), thoracic (T1-T6) and lumbar (L1-L2) spinal cord. Some of these regions have previously been shown to have functionally specialized roles^77,78^ and very likely contain dedicated neuron types that are not present in our atlas. Furthermore, although we identify a large number of projection neurons, we are only detecting the spinal cord output cells that take up rgAAV capsid payloads. However, rgAAV viral capsid has been described to have capsid tropism effects in some central neurons^79,80^ which may also be the case for the spinal cord cells. Moreover, several of the brain structures we targeted for retrograde tracing (e.g. LPBN, CVLM) are localized near major ascending fiber pathways which may have taken up viral tracers without any direct innervation potentially inflating projection collateralization estimates at these structures. This issue has recently been better characterized with passing fiber uptake of rgAAV tracers ranging from negligible to considerable depending on the fiber pathway^79,81^. Furthermore, viral spread to adjacent brain regions and along intracranial injection paths (cerebellum/LBPN, brainstem structures) may also confound interpreting wiring patterns. Thus, weaker connections in the connectivity matrix as well as connectivity in structures near major ascending fiber tracts should be interpreted with caution. Finally, although most SPN categories — including DSPNs, LSPNs, CC-SPNs, PSPNs, and L10-SPNs — contain transcriptomically discernible neuron types, some DP-SPNs are less distinct from transcriptomically similar non-projecting neurons. Thus, future work with carefully re-designed spatial transcriptomic probe sets will likely refine the composition of DP-SPN category projection neurons.

### Outlook

The atlas, online portal, and accompanying nomenclature establish a shared framework for the molecular, anatomical, and functional dissection of the mammalian spinal cord. Linking the >500 neuron types described here to physiological tuning, circuit connectivity, developmental origin, and behavioral output—and ultimately to homologs in human spinal cord—will require a coordinated community effort spanning genetics, viral tools, *in vivo* imaging, and behavior. The framework we provide is a step toward that goal.

## Experimental Procedures

### Animals

All animal care and experimental procedures were carried out in accordance with the US NIH guidance for the care and use of laboratory animals and approved by the University of Texas Southwestern Medical Center Animal Care and Use Committee (protocol #2022-103301). Mice used for multiomic and spatial transcriptomic profiling of the spinal cord were 8-week-old C57Bl/6J male and female mice. For retrograde viral tracing, 8-week-old male and female mice received intracranial viral injections followed by 9-11 day incubation period and tissue collection. All mice were single-housed for three days prior to tissue collection to avoid fighting induced transcriptional artifacts. Mice were housed in temperature- and humidity-controlled rooms with a 12h light/dark cycle and unrestricted access to chow and water.

### Spinal Cord Sample Processing for 10x Genomics Multiomic profiling

Tissue collection and processing into profiling ready nuclear suspensions was performed as previously described^82^. Briefly, 8-week-old male and female C57Bl/6J mice were anesthetized with isoflurane in a drip chamber and animals were decapitated. Spinal cord was rapidly surgically extracted and transferred to ice-cold carbogenated (95% O_2_ and 5% CO_2_) NMDG-Hepes-ACSF (93 mM NMDG, 2.5 mM KCl, 1.2 mM NaH_2_PO_4_, 30 mM NaHCO_3_, 20 mM HEPES, 25 mM glucose, 10 mM MgSO_4_, 1 mM CaCl_2_, 1 mM kynurenic-acid Na salt, 5 mM Na-ascorbate, 2 mM thiourea and 3 mM Na-pyruvate, pH adjusted to 7.4). Spinal cord tissue was separately isolated from cervical (C3-C8), thoracic (T7-13), lumbar (L1-5) and sacral (S1-S4) segments. Samples were immediately transferred and dissociated using a Dounce tissue grinder (Kimble #885300-0002) in lysis buffer (0.32 M Sucrose, 10 mM Tris-HCl pH=7.4, 3 mM MgCl_2_, 10 mM NaCl, 0.50% BSA, 0.1% Triton-X-100 and 0.1U/µl Protector RNAse Inhibitor (Roche #RNAINH-RO) in Ultrapure water (Invitrogen #10-977-023), 1 mM DTT, 1x EDTA free Protease Inhibitor (Roche #11873580001)) under RNase-free conditions. The homogenate was filtered through a 40 µm cell strainer (Fisher #NC1789427) and layered onto a 12 mL Sucrose Cushion solution (1M sucrose, 10 mM Tris-HCl pH=7.4, 3 mM MgCl_2_, 10 mM NaCl, 0.50% BSA, 0.10% Triton-X-100 and 0.1U/µl Protector RNAse Inhibitor (Roche #RNAINH-RO) in Ultrapure water (Invitrogen #10-977-023), 1 mM DTT, 1x EDTA free Protease Inhibitor (Roche #11873580001)) in 50 mL Oakridge tubes (Thermo Fisher #0556214D). Samples were centrifuged at 3,200g for 20 minutes at 4°C on a spin-out rotor. The supernatant was rapidly decanted and the nuclear pellet was resuspended in nuclear buffer (0.32 M Sucrose, 10 mM Tris-HCl pH=7.4, 3 mM MgCl_2_, 10 mM NaCl, 0.50% BSA, and 0.1U/µl Protector RNAse Inhibitor (Roche #RNAINH-RO) in Ultrapure water (Invitrogen #10-977-023), 1 mM DTT, 1x EDTA free Protease Inhibitor (Roche #11873580001)). To isolate neuronal nuclei, nuclear suspensions were incubated with an anti-NeuN antibody (Biolegend #834501) custom conjugated to Alexa Fluor 647 (Molecular Probes #A30009, 1:100 diolution) and 7-AAD (10µg/ml) for 15 minutes at 4°C in the dark. Nuclei were spinned down at 500g for 5 min and resuspended in nuclear buffer twice to wash out stains followed by Fluorescence-activated cell sorting (FACS) on either FACS Aria II or FACS Aria Fusion cell sorters at low sorting speeds (4-5) after compensating for spectral overlap with single stained samples. 100 000 – 120 000 7-AAD/anti-NeuN double-positive neuronal nuclei were sorted into the nuclear buffer. Nuclear suspensions were concentrated with CUTANA Concanavalin A beads (Epicypher # 21-1401) according to manufacturer’s protocol and resuspended in 15 microL of 1x Nucleus Buffer (10x Genomics, part of #1000283) yielding a nucleus concentration of ∼4000 nuclei/µl. The nuclear suspensions were profiled in three parallel technical replicates/wells on the 10x Genomics Chromium iX instrument following the manufacturer’s protocol (10x Genomics, #1000283) yielding snRNA-seq and snATAC-seq Illumina sequencing libraries. The snMultiome libraries were sequenced on the Novaseq X 10B flow cells at 100 000+ mean raw reads/cell for both snRNA-seq and snATAC-seq libraries. Reads from snMultiome were mapped to the 10x Genomics mouse transcriptomic reference (mm10-2020-A-2.0.0) using Cell Ranger Arc 2.0.2 with default parameters resulting in gene-cell and peak-cell matrices.

### Pre-processing and Analysis of Single-Nucleus RNA-Sequencing Data

Ambient mRNA background was removed from gene-cell matrices with Cellbender (parameters: --expected cells <CELL Ranger Arc cell count estimate>, --total-droplets-included 1.75*expected_cells, --fpr 0.01, epochs 150, --exclude-feature-types Peaks, --projected-ambient-count-threshold 0.1, zdim 40). Resulting gene expression data was processed with the Seurat v5.3.0 R package. Low quality nuclei and partially lysed cells were filtered out by thresholding out nuclei with less than 750 unique transcripts (UMI) and more than 1% of mitochondrial reads. Non-neuronal cells and neuron-non neuron doublets were removed from the data based on expression of canonical marker genes for oligodendrocytes, astrocytes, ependymal cells and other non-neural cell classes based on previously reported cell class markers^46^. Neuron-neuron doublets were identified with scDblFinder^83^ and clusters comprising high doublet probability cells with no unique markers were removed. The resulting QC passed data was separated into glutamatergic, GABAergic and cholinergic neuron datasets for independent analysis by high granularity Louvain clustering and separating clusters based on expression of canonical glutamatergic (*Slc17a6*), GABAergic (*Gad1, Slc32a1*) and cholinergic markers (*Chat*). Neurons in clusters containing mixed but non-overlapping GABAergic and glutamatergic marker expression were disaggregated into respective datasets by isolating glutamatergic neurons (Slc17a6 count>Gad1+Slc32a1 count) from GABAergic neurons (Slc17a6 count<Gad1+Slc32a1 count) and separating to respective neurotransmitter specific datasets. We did not observe reliably glutamatergic and GABAergic marker co-expressing neurons across Multiome and Xenium modalities that were not doublets/segmentation artifacts in follow-up analysis. However, cholinergic neurons often co-expressed GABAergic or glutamatergic markers, which has previously been well documented^13,14^. Therefore, cholinergic neurons with only one exception (CHOL-GABA-Crnde) were separated into the cholinergic dataset regardless of their glutamatergic/GABAergic marker expression.

We performed crude unsupervised Louvain clustering separately on glutamatergic, GABAergic and cholinergic neurons to yield transcriptomic groups broadly corresponding to transcriptomic families^7,46^/groups^45^ in recent studies or neuron types in earlier^1,4^ atlases (**Figure S1A**). Transcriptomic groups were named based on their neurotransmitter (GLUT/GABA/CHOL) and an evolutionarily conserved broad marker consolidated across the rodent and primate lineages with a companion study^45^. Furthermore, each transcriptomic group was individually subclustered into neuron types using unsupervised Louvain clustering as implemented by the Seurat R package. Briefly, we identified between 500-1250 differentially expressed genes for each group with the “vst” algorithm as implemented by the FindMarkerGenes function on log-transformed [ln(1+(gene count × 10,000)/total transcript count per cell)] data. Gene expression values were z-scored across cells and dimensionality reduction was performed with principal component analysis. We used principal components accounting for the most variance (as determined by the ‘elbow’ in the Scree plot) ranging from 8 to 24 PCs as input to clustering analysis. We compiled the shared nearest-neighbour graph and identified transcriptomic clusters by optimizing the graph modularity function with the Louvain algorithm as implemented by Seurat::FindNeighbours and Seurat::FindClusters (resolution parameter ranging from 0.1 to 1.5). We determined the optimal clustering resolution by evaluating cluster stability with the ClusterTree R package choosing the minimal resolution yielding stable clusters across largest span of resolution values that yields transcriptomic groups with more than 10 selective genes for most clusters within the group. Individual neuron types were named based on their neurotransmitter profile (GLUT/GABA/CHOL), broad marker from group name and finally fine marker uniquely defining neuron type with the group. Final neuron type nomenclature and naming was consolidated by cross-referencing marker gene expression with Xenium analysis prioritizing fine marker choice for neuron type naming that would be detectable across both modalities.

### Pre-processing and Analysis of snATAC-seq Data

snATAC seq data from the multiomic experimements were processed using Seurat (v5.3.0) R package and Signac (v1.16.0). Peaks were called per pre-defined neuron groups on the aggregated fragments file from 4 samples ("AC04A", "AL03A", "AS03A", "AT01B") using MACS2 (v2.2.7) via Signac’s CallPeaks() function. A peak-by-nucleus matrix was constructed using FeatureMatrix(). Peaks below the 25th percentile of detection frequency were excluded. Accessibility counts were normalized by term-frequency inverse-document-frequency (TF-IDF) transformation and dimensionality reduction was performed by LSI (via SVD). The first LSI component, which correlates with read depth, was excluded from subsequent analyses.

Differentially accessible peaks per cluster were identified using logistic regression (FindMarkers(), test.use = "LR") with sequencing depth (nCount_peaks_by_cluster) included as a latent variable, retaining only positive associations (peaks more accessible in the focal cluster) with a minimum detection fraction of 5% and log-fold-change threshold of 1.

### Analysis of Neuron Type Rostro-Caudal Prevalence

Rostro-caudal prevalence of neuron types was estimated by first anatomically balancing the sampling from individual multiomic datasets (glutamatergic, GABAergic and cholinergic neurons) between the four rostro-caudal anatomic regions (cervical, thoracic, lumbar, sacral) downsampling all data to least abundant anatomical region to avoid sampling depth based false positives. We used convergence of two statistical tests to call high-confidence rostro-caudally varying neuron types.

To identify cell types with statistically significant compositional differences across spinal cord anatomical regions, we applied two complementary statistical frameworks: DESeq2-based negative binomial modeling and beta regression on proportional data. A cell type was considered to show significant regional variation only if it reached statistical significance in both models.

Analyses were performed on a cell count matrix tabulating the number of cells per cell type per biological sample (mouse), constructed from a dataset balanced by both anatomical region and sex. Sample identifiers encoded both sex and spinal cord region of origin.

DESeq2 Likelihood Ratio Test: Raw cell type counts were analyzed using DESeq2 (v1.44.0) to test for regional effects while controlling for sex. A DESeqDataSet object was constructed with the design formula ∼ sex + region, where region levels included cervical (AC), thoracic (AT), sacral (AS), and lumbar (AL). A likelihood ratio test (LRT) was used to compare the full model (∼ sex + region) against the reduced model (∼ sex), thereby isolating the contribution of anatomical region to cell type abundance variation. Size factor normalization and dispersion estimation were performed using default DESeq2 parameters. For each pairwise regional comparison, r:esults were extracted using explicit contrasts and a cell type was considered differentially abundant between a given pair of regions if the Benjamini-Hochberg (BH)-adjusted p-value was < 0.05 and the absolute log₂ fold change exceeded log₂(3) (i.e., a minimum 3-fold difference). All pairwise combinations among the four regions were evaluated.

Beta Regression: To complement the count-based analysis, we modeled cell type proportions using beta regression (betareg v3.2.4), which is appropriate for response variables bounded on the interval (0, 1). For each biological sample, raw counts were converted to proportions such that all cell type fractions within a sample summed to 1. Proportions were transformed using the formula (y × (n − 1) + 0.5) / n to avoid boundary values of exactly 0 or 1, where n is the number of samples. The beta regression model was specified as proportion ∼ region + sex, with lumbar (AL) as the reference region. For each cell type, regional coefficients (AC, AT, AS relative to AL) were extracted and BH correction was applied jointly across all cell types and all model terms simultaneously. A cell type was considered significant if at least one regional coefficient had a BH-adjusted p-value < 0.05.

### Viral constructs

The following viruses for retrograde viral tracing were obtained from Addgene: rgAAV-CAG-tdTomato (Addgene, #59462-AAVrg), rgAAV-CAG-GFP (Addgene, #37825-AAVrg). Furthermore, we obtained a 711 bp mNeonGreen synthesized coding sequence from Twist Bioscience (sequence from Addgene #99133) and cloned into the pAAV-CAG-fluorophore backbone (Addgene #59462). The resulting AAV genome was packaged into a rgAAV capsid at Neurotools yielding a third retrograde tracing virus rgAAV-CAG-mNeonGreen. The coding sequences of the used retrograde viral tracers were orthogonal enabling multiplexed independent quantification on the Xenium platform.

### Stereotactic surgery for retrograde tracing

Retrograde viral tracers were introduced to major spinal cord output brain structures via intracranial surgeries as previously described^60^. Briefly, animals were anaesthetized with a mixture of ketamine (1 mg/ml) and xylazine (10 mg/ml) in isotonic saline, injected intraperitoneally (i.p.) at a dose of 10 μl/g body weight. Meloxicam was subcutaneously administered at 2 mg/kg body weight. The mouse was then placed in a stereotaxic apparatus (Narishige #SR-5M-HT) on a heating pad at 37 °C. The mouse skull was exposed by an incision of the scalp followed by application of the topical analgesic lidocaine (2.5 mg/ml). Small craniotomies, less than 1 mm, were made using a hand drill at the regions of interest. Virus injections were performed with pulled glass pipettes using a microprocessor-controlled injection system (Nanoliter 2020, World Precision Instruments) at 100 nl/min. The following brain regions were targeted for retrograde tracing with respective virus preparations and volumes: mediodorsal thalamus (MD; AP: -1580, ML: +/- 500, DV: +3250; rgAAV-mNeonGreen, 200 nL), ventral posterolateral nucleus of the thalamus (VPL; AP: -1400, ML: +/- 1900, DV: +3550; rgAAV-mNeonGreen, 200 nL), posterior triangular thalamic nucleus (PoT; AP: +3100, ML: +/- 1600, DV: +3650; rgAAV-tdTomato, 200 nL), superior colliculus (SCol; AP: +4700, ML: +/- 500, DV: 1500; rgAAV-mNeonGreen, 125 nL), periaqueductal grey (PAG; AP: −5000, ML: +/-550, DV: +2700; rgAAV-tdTomato, 150-200 nL), lateral parabrachial nucleus (LPBN; AP: −5300, ML: +/- 1275, DV: +3625; rgAAV-tdTomato, 175 nL), cerebellum (CEB; AP: +6200, ML: +/- 1000, DV: 2700 & 3000; rgAAV-mGFP, 175 nL per injection site), caudal ventrolateral medulla (CVLM; AP: −7600, ML: +/-1325, DV: +5850; rgAAV-GFP, 150 nL), dorsal column nuclei (DCN; AP: -7800, ML: +/- 300, DV:+5000; rgAAV-GFP, 100 nL). Following intracranial injections, the scalp incision was closed with GLUture topical tissue adhesive (Abbott Laboratories #032046). After the surgery, all mice received a subcutaneous injection of buprenorphine SR at 1 mg/kg body weight and were placed in a clean cage on a heating pad overnight to recover and were then housed in the animal facility. A total of 4 animals (2 male, 2 female) were injected and profiled per spinal cord projection target site. 9-11 days after intracranial surgeries, animals were euthanized, spinal cords were collected and embedded in OCT for spatial transcriptomic profiling. Brains and spinal cord regions outside collection segments were fixed in 4% PFA overnight, sectioned on Leica VT1000S vibratome followed by DAPI staining and high-speed camera imaging on Leica Stellaris 8 to validate virus delivery to target brain regions.

### Spinal Cord Sample Processing for 10x Genomics Xenium Spatial Transcriptomic Profiling

8-10-week-old male and female C57Bl/6J mice were anesthetized with isoflurane in a drip chamber and animals were decapitated. Spinal cord was rapidly surgically extracted and transferred to ice-cold carbogenated (95% O_2_ and 5% CO_2_) NMDG-Hepes-ACSF (93 mM NMDG, 2.5 mM KCl, 1.2 mM NaH_2_PO_4_, 30 mM NaHCO_3_, 20 mM HEPES, 25 mM glucose, 10 mM MgSO_4_, 1 mM CaCl_2_, 1 mM kynurenic-acid Na salt, 5 mM Na-ascorbate, 2 mM thiourea and 3 mM Na-pyruvate, pH adjusted to 7.4). Spinal cord tissue was separately isolated from cervical (C4-8), thoracic (T8-13), lumbar (L4-5) and sacral (S1-2) segments of wild-type C57Bl/6J mice and embedded in Tissue-Tek O.C.T. compound using a liquid nitrogen-isopentane bath for snap freezing and stored at -80°C. 10 µm thick cryosections were cut on Leica CM3050S Cryotome and mounted onto Xenium slides (10x Genomics, #3000941). Fixation was performed with 4% paraformaldehyde (PFA), freshly prepared from a 16% stock solution (15710, Electron Microscopy Sciences) at room temperature for 30 minutes, followed by permeabilization with pre-chilled 70% methanol (34860, Millipore Sigma) on ice for 1 hour, according to the Xenium In Situ for Fresh Frozen Tissues – Fixation & Permeabilization protocol (CG000581). All steps were carried out under RNase-free conditions. Probe hybridization and cell segmentation were performed following the manufacturer’s protocol Xenium In Situ Gene Expression with Cell Segmentation Staining (CG000749). Profiling ready tissues were processed on the Xenium Analyzer and resulting Xenium objects were generated by Xenium Analyzer Software 3.3.0.1.

### Design, Preprocessing and Analysis of Xenium Spatial Transcriptomic Data

We designed a stand-alone custom gene panel for Xenium v1 chemistry to detect the expression of 480 genes including key marker genes for spinal cord transcriptomic groups and cell/neuron types based on our multiomic profiling as well as genetic markers from spinal cord literature (**Table S2**). Xenium spatial transcriptomic data was processed and analyzed with the Seurat v5.3.0 R package with added visualization functions from Xenium Explorer v4.0. We removed low quality segmentation products by excluding cells with fewer than 20 transcripts. Cells were annotated with anatomical (rostro-caudal region), experimental (trace vs. no trace and origin of tracing signal) and sex identity. Sex identity of spinal cord sections was validated by Y-linked *Ddx3y* gene expression. Tracing and Y-linked sex marker genes were removed from the gene-cell matrix and stored in metadata to avoid biasing cell-type identification. Neuronal data was extracted from full spatial transcriptomic data and neuron-non-neuron segmentation doublets were excluded based on canonical marker expression (Rbfox3, Slc17a6, Slc32a1 and non-neuronal cell class markers). Neuronal data was segregated into glutamatergic, GABAergic and cholinergic datasets based on expression of canonical glutamatergic (*Slc17a6*), GABAergic (*Slc32a1*) and cholinergic markers (*Chat*). Xenium data was normalized to [ln(1+(gene count × median(RNA count))/total transcript count per cell)]. Group level identities were transferred from snRNA-seq datasets by using CCA integration based label transfer^84^. Briefly, snRNA-seq and Xenium spatial transcriptomic data was integrated with CCA separately for each neurotransmitter dataset. We performed granular Louvain clustering on resulting integrated data (474 shared genes, 30 PCs, clustering resolution 3.5) and majority snRNA-seq labels in each cluster were transferred to the Xenium neurons. Resulting transcriptomic group labels were manually curated to adjust for label transfer errors based on group level markers (**Figure S1B-D**). Neuron-neuron segmentation doublets were removed based on group level marker overlap as well as confirming anatomical feasibility of doublets based on anatomical annotation of groups (**Figure S4, S5**). In case of ambiguities, candidate doublets were directly visually inspected in Xenium Explorer to validate or rule out doublet status and exclude doublet clusters. For each individual glutamatergic and GABAergic group, we performed unsupervised Louvain clustering (476/480 genes excluding *Ddx3y* and tracer genes; 8-27 PCs, resolution: 0.1-2.5) followed by CCA based label transfer from multiomic data to Xenium spatial transcriptomic data to assign first pass neuron type identity. The resulting Xenium neuron type identities were refined and curated based on presence or absence of shared marker genes. Xenium clusters with low label transfer prediction scores and no clear corresponding neuron type in multiome data were assigned a new identity. For the cholinergic neuron dataset, we performed de-novo Leiden clustering and cell-type identification for all 5 groups as multiomic cholinergic dataset was underpowered/sampled. Corresponding cholinergic neuron types were determined by targeted marker gene expression analysis. Finally, resulting neuronal nomenclatures and neuron type names were harmonized across multiomic and Xenium datasets prioritizing fine marker name components from genes clearly detected across the two profiling chemistries.

### Identification of Sex Specific Neuron Types in Spatial Transcriptomic Data

To identify cell types with statistically significant abundance differences between sexes in the Xenium spatial transcriptomics dataset, we applied DESeq2 (v1.44.0) likelihood ratio testing on cell type count data. The input count matrix tabulated the number of cells per cell type per biological sample.

A DESeqDataSet object was constructed with the design formula ∼ sex. A likelihood ratio test (LRT) was then used to compare this full model against an intercept-only reduced model (∼ 1), directly testing whether sex explains a significant proportion of variance in cell type abundance. Size factor normalization and dispersion estimation were performed using default DESeq2 parameters. Multiple testing correction was applied automatically using the Benjamini-Hochberg (BH) method to control the false discovery rate (FDR). A cell type was considered to show significant sex-dependent abundance differences if the BH-adjusted p-value was < 0.05.

### Identifying spinal projection neurons from retrograde tracing data

Spinal projection neurons (SPNs) were identified from spatial transcriptomic data using raw retrograde tracer counts (tdTomato, GFP, or mNeonGreen) from four animals per spinal cord target structure (two male and two female). Because retrograde tracing introduces technical artifacts that are absent in non-traced animals — including segmentation errors and false-positive signal arising from labeled axons of passage — count distributions from non-traced animals could not provide a valid noise reference. We therefore used GABAergic neurons from the same traced animals as an internal reference, on the basis that these cells are subject to the same sources of technical noise as candidate populations while contributing minimally to ascending projections.

We note that a small number of inhibitory ascending projection neurons have been reported^41,85^. Using GABAergic neurons as the reference therefore biases the analysis toward false negatives and yields conservative SPN estimates.

For each tracer, we tested each candidate neuron type against this internal reference using a permutation test. Tracer count values from cells of a given neuron type were assigned the "candidate" label, and tracer count values from all GABAergic neurons in the traced experiments were assigned the "control" label. The test statistic was defined as the difference between the mean candidate count and the mean control count. An empirical null distribution was generated by randomly permuting the candidate/control labels 100,000 times, and the empirical p-value was computed as the fraction of permuted statistics meeting or exceeding the observed value. P-values were corrected for multiple comparisons using the Bonferroni method across 1,014 tests spanning the three tracer experiments.

To exclude neuron types with insufficient tracer signal, we required each retained SPN type to contain at least 5 cells with >5 tracer transcripts for the GFP and tdTomato experiments, and at least 4 cells with >5 tracer transcripts for the mNeonGreen experiments; the lower count threshold for mNeonGreen reflects the reduced labeling efficiency of the mNeonGreen rgAAV relative to the GFP and tdTomato rgAAV tracers. We further excluded any cluster in which fewer than 1% of cells exceeded these per-cell thresholds.

### Spatial registration of neuron types to reference atlas

Cell centroids from spatial transcriptomic data were registered to a common coordinate system from anatomic reference segments from either a cervical (C5), thoracic (T12), lumbar (L5), or sacral (S1) spinal cord obtained from the Allen Institute Mouse Spinal Cord Atlas (https://mousespinal.brain-map.org/) with a custom code. Briefly, spatial coordinates for spinal cord cells and transcripts were loaded from a Seurat object. We defined 22 anchor points along the spinal cord gray matter boundary capturing key anatomical landmarks both on the reference contours as well as the spinal cord datasets. Gray matter in individual spinal cords was visualized by the gene expression of neuronal markers (*Slc17a6, Slc32a1, Chat*) that served as guides for manually defining anchor points for registration. A spatial transformation model was estimated by fitting a thin plate spline surface (TPS) to source (spinal cord) and target (reference atlas) anchor points and used to transform cell centroid data to common reference atlas coordinates using spatial data input/output, modeling and manipulation functions as implemented in imager, fields, sp and morpho R libraries. We visualized cell-type-specific anatomical locations by aggregating cell location data across 20 (10 male and 10 female) anatomical sections.

### Label transfer for spinal projection neuron datasets

We used common correlation analysis (CCA) based label transfer as implemented in Seurat v5.3.0 R package to identify corresponding neuron types in three published single-nucleus spinal projection neuron datasets including the current study (Cano-Gomez et al., Xenium v1 chemistry), Phox2a lineage lumbar SPNs (Bell et al. 2024, snRNA-seq Smart-seq2 chemistry) and cervical retro-seq identified SPNs (Lin et al. 2026, snRNA-seq 10x Genomics v3 chemistry). To overcome technical differences in single-cell profiling chemistries, the label transfer parameters were adjusted for each pair-wise label transfer. Briefly, to transfer Cano-Gomez et al. lumbar SPN labels to Phox2a+ lumbar SPNs (Bell et al. 2024) we identified overlapping genes between 476 Xenium genes and 15k Phox2a most variable features (285 genes) identified by the “vst” algorithm. Cano-Gomez et al. labels were transferred using shared CCA embedding based on transfer anchors established on first 14 principal components chosen based on plateau value in the scree plot. Low confidence label transfer events were filtered by including cells with prediction scores above 0.4 in final analysis. Furthermore, Cano-Gomez et al. cervical SPN labels were transferred to Lin et al. cervical SPNs by first identifying overlapping genes between 476 Xenium genes and 5k most variable cervical SPN genes (300 genes) identified by the “vst” algorithm. Cano-Gomez et al. labels were transferred using shared CCA embedding based on transfer anchors established on first 14 principal components. Low confidence label transfer events were filtered by including cells with prediction scores above 0.6 in final analysis. Finally, Lin et al. cervical SPN labels were transferred to the lumbar Phox2a cells by first identifying overlapping genes between the 4000 most variable genes (972 genes) from both datasets as identified by the “vst” algorithm. Lin et al. labels were transferred using shared CCA embedding based on transfer anchors established on first 16 principal components. Low confidence label transfer events were filtered by including cells with prediction scores above 0.6 in final analysis

### Generation of cell-type-to-phenotype maps

We used the spinal projection neuron atlas data generated by the Xenium spatial transcriptomic platform to generate the cell-type-to-phenotype map of spinal projection neurons. The purpose was to provide a cell-type-resolved reference to contextualize functional SPN level interventions that yield supraspinal somatosensory behavioral phenotypes. To this end, we included genetic markers from all studies demonstrating supraspinal somatosensory behavioral phenotypes as a result of SPN specific functional intervention (opto- or chemogenetic activation/silencing and SPN targeted genetic perturbations) at the time of Xenium gene panel design. We used broad phenotypic categories of “pain”, “itch”, “touch” and “cold sensation” to aggregate behavioral experiments testing a variety of behavioral phenotypes under these umbrella terms. For example, “chemical itch“ and “mechanical itch” behaviors were aggregated under “itch phenotype” and different supraspinal pain assay phenotypes were aggregated under “pain phenotype”. We discretized the marker gene expression in the SPN reference atlas by non-zero quantile thresholding^86^. Briefly, we calculated bulk-average marker gene expression for each SPN type using the AverageExpression() function in Seurat v.5.3.0. Cell-type level average expression estimates were thresholded by median expression value of the gene across all non-zero expressing cells in the atlas. The resulting 1 or 0 expression estimates threshold out cells with very low and inconsistent expression of marker genes as well as occasional segmentation error derived background gene expression noise. Resulting discrete marker gene expression estimates were plotted across studies to reveal underlying targeted neuron types for specific manipulations. Some of the studies involved intersectional approaches where the effect of the genetic manipulation was defined by the intersection of a Cre and FlpO genetic reagents with the latter restricting effector gene expression to a particular developmental lineage (e.g. Lbx1-FlpO). As the adult expression of the developmental markers may not represent not reflect all developmentally impacted cells, we used the full Cre driver line expression to estimate the population of manipulated cells to skew more conservative about precision of the final maps.

## Resource Availability

### Lead contact

Further information and requests for resources and reagents should be directed to and will be fulfilled by the lead contact, Allan-Hermann Pool

### Materials availability

The pAAV-CAG-mNeonGreen plasmid generated in this study is available from the lead contact upon request. The 480-gene custom Xenium panel gene list is provided in **Table S1**, with panel design details available from the lead contact. All other reagents are commercially available or were obtained from the sources indicated in the key resources table.

### Data and code availability

Raw and summary snMultiome data is available at Gene Expression Omnibus (accession #GSE318731).

Xenium Spatial Transcriptomic data is available at Brain Imaging Library (doi: www.doi.org/10.35077/g.1196).

Code for data analysis in this manuscript is available at https://github.com/PoolLab/cano-gomez_et_al_2026.

Code for registering spatial transcriptomic data to reference atlas: https://github.com/isst001/spatial-image-registration.git

## Supplemental Information

### Author Contributions

A-H.P. and H.C.L. conceived the project. A-H.P., H.C.L., and S.I., designed the experiments.

S.I. developed single-nucleus and spatial transcriptomic tissue processing workflows and established image registration analysis for spatial mapping. L.C.G. performed multiomic profiling of spinal cord tissue with assistance from H.P. L.C.G. and S.I. performed spatial transcriptomic profiling of spinal cord tissue, with assistance from H.P. S.I. first characterized transcriptomic SPN types in the lumbar spinal cord and led the analysis and characterization of SPN diversity. H.P., A-H.P. and E.I. analyzed the data. H.P. designed and implemented the online data analysis portal. H.C.L. and A-H.P. acquired funding. A-H.P., H.P. and S.I. wrote the manuscript. All authors edited the manuscript. A-H.P. supervised the project.

## Supporting information

Table S1: Multiomic Adult Mouse Spinal Cord Nomenclature.

Table S2: Xenium gene probe set used for spatial transcripomic profiling.

Table S3: Spatial transcriptomic adult mouse spinal cord nomenclature.

## Acknowledgements

We thank members of the Pool lab and Dr. Seungwon Choi for feedback on the manuscript. We thank Michael Ortiz and the Moody Flow Cytometry Core at UTSW for advice and execution of flow cytometry. We thank members of the BRAIN Initiative Cell Atlas Network (BICAN) Spinal Cord Working Group including Dr. Matthew Scmitz, Dr. Trygve E. Bakken, Dr. Michael J. Leone, Dr. Nelson Johansen for discussions on mammalian spinal cord nomenclature consolidation.

A-H.P. is supported by UT Southwestern McDermott Endowed Scholars Award, Rita Allen Scholar Award in Pain, McKnight Foundation Neurobiology of Brain Disorders Award, and BICAN R01MH134799 from NIMH. S.I. is supported by JSPS Overseas Research Fellowship. H.C.L. is supported by R01MH134799 from NIMH.

## Supplemental Information

Document S1: Extended Data Figures S1-S11.

Interactive portal for spinal cord cell-type, anatomy and projectome analysis: https://spinal-cord-explorer.shinyapps.io/mouse/

Table S1: Multiomic adult mouse spinal cord nomenclature.

Table S2: Xenium gene probe set used for spatial transcripomic profiling.

Table S3: Spatial transcriptomic adult mouse spinal cord nomenclature.

---

**Extended Data Figure 1:**
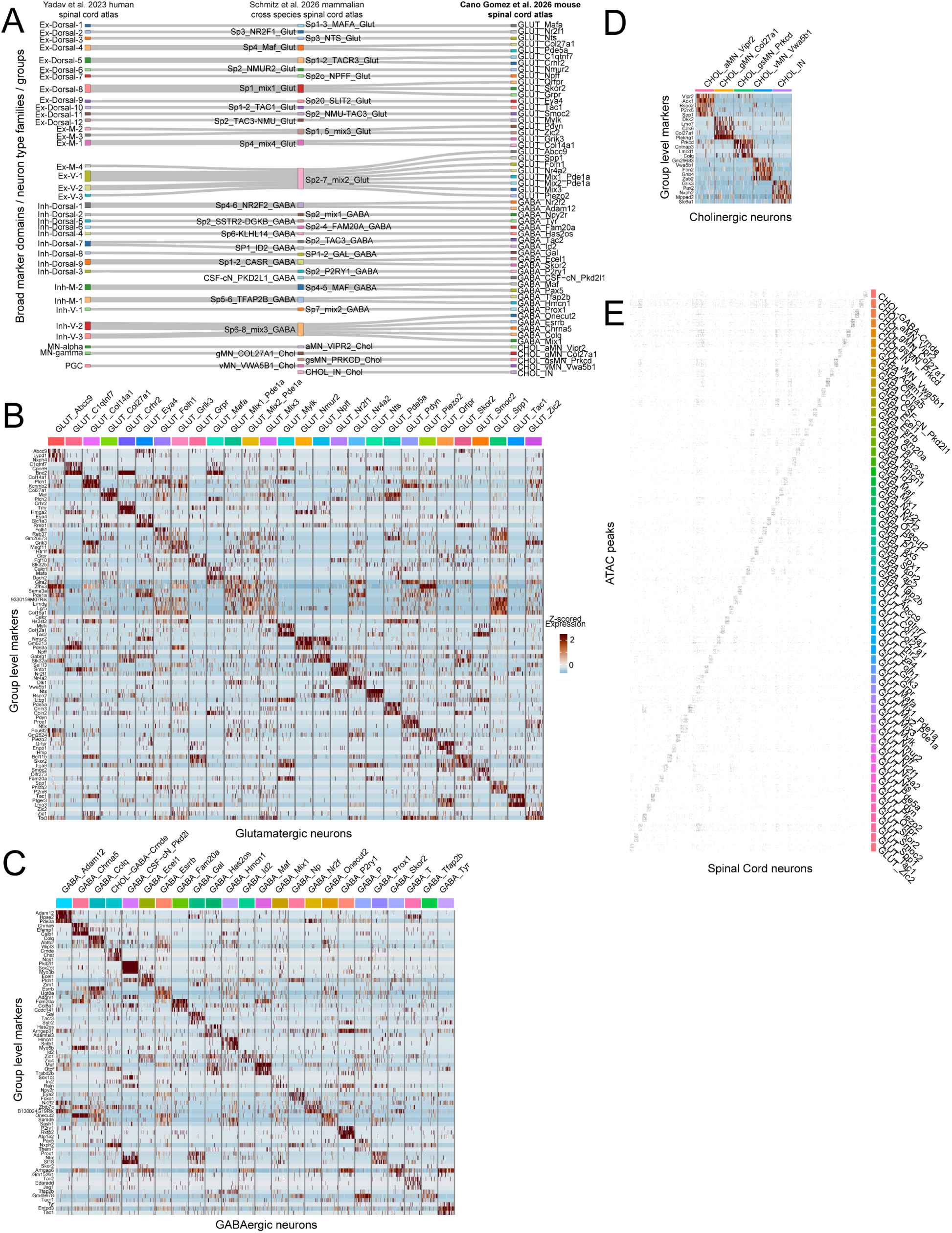
Molecular characterization of spinal cord neuronal diversity with multiomic profiling. (A) Group level neuronal nomenclature and corresponding neuron types/transcriptomic groups in a companion pan-mammalian spinal cord study (Schmitz et al. 2026) and a recent human spinal cord atlas (Yadav et al. 2023). (B) Heatmap of glutamatergic group level markers in the adult mouse spinal cord. (C) Heatmap of GABAergic group level markers in the adult mouse spinal cord. (D) Heatmap of cholinergic group level markers in the adult mouse spinal cord. (E). Heatmap of group level open chromatin regions in adult mouse spinal cord.

**Extended Data Figure 2:**
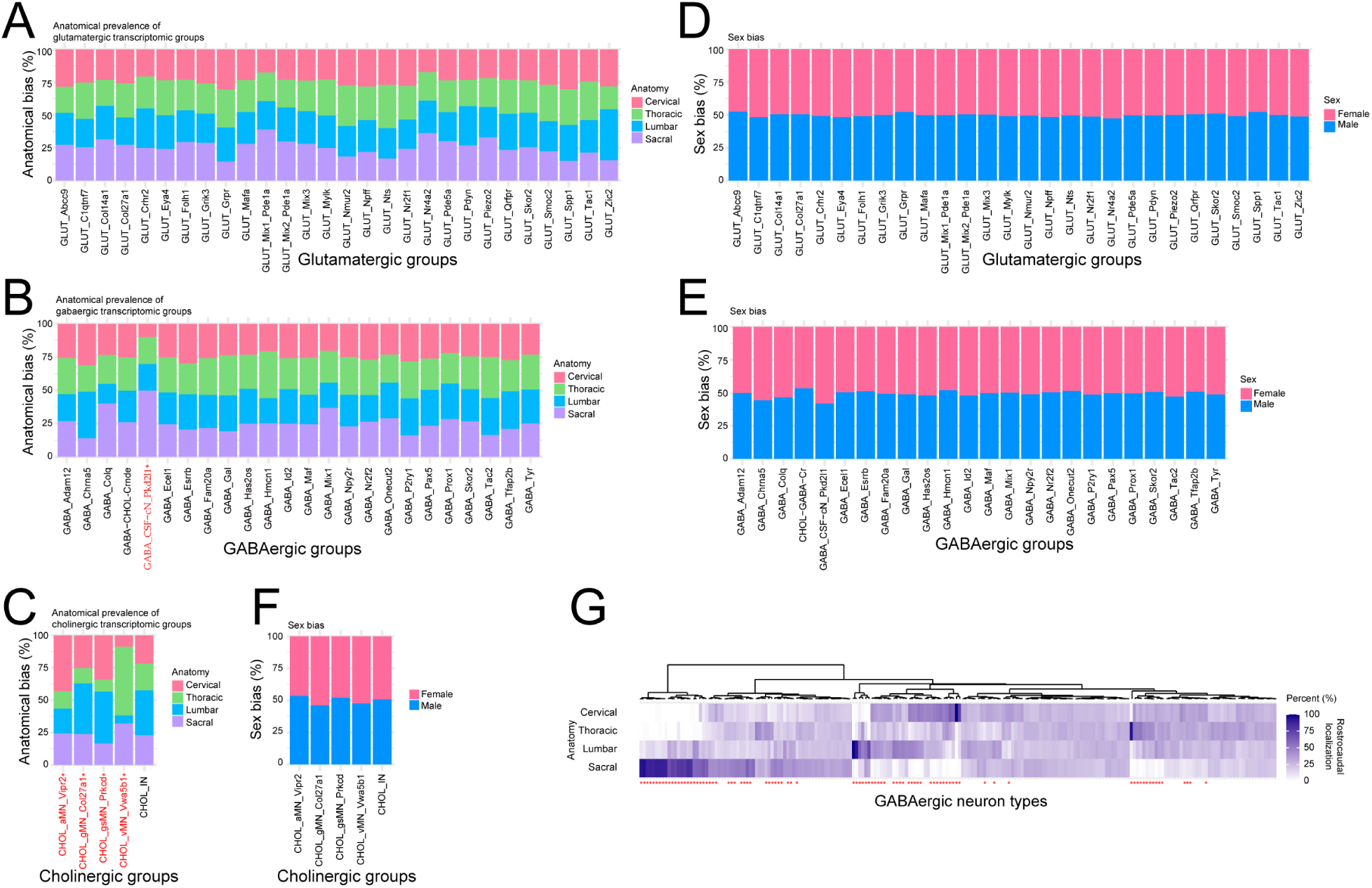
Group and neuron type level rostro-caudal distribution and sex biases for multiomic spinal cord dataset. (A) Rostro-caudal distribution of glutamatergic groups in a regionally balanced sample (n= 86 740 cells). (B). Rostro-caudal distribution of GABAergic groups in a regionally balanced sample (n= 66 116 cells). Groups marked by red asterisk/text are rostro-caudally significantly specialized (three-fold enrichment between at least two rostro-caudal regions, adjusted p<0.05 DeSeq2 and adjusted p<0.05 with beta regression). (C) Rostro-caudal distribution of cholinergic groups in a regionally balanced sample (n=2012 cells). (D) Glutamatergic group level male/female bias from a sex balanced sample (n=87 830 cells). (E) GABAergic group level male/female bias from a sex balanced sample (n=70 558 cells). (F) Cholinergic group level male/female bias from a sex balanced sample (n=2524 cells). (G) Rostro-caudal distribution of GABAergic neuron types in the mouse spinal cord. Neuron types marked in red are rostro-caudally significantly specialized (three-fold enrichment between at least two rostro-caudal regions, adjusted p<0.05 DeSeq2 and adjusted p<0.05 with beta regression, see **Table S1** for gabaergic neuron identity labels).

**Extended Data Figure 3:**
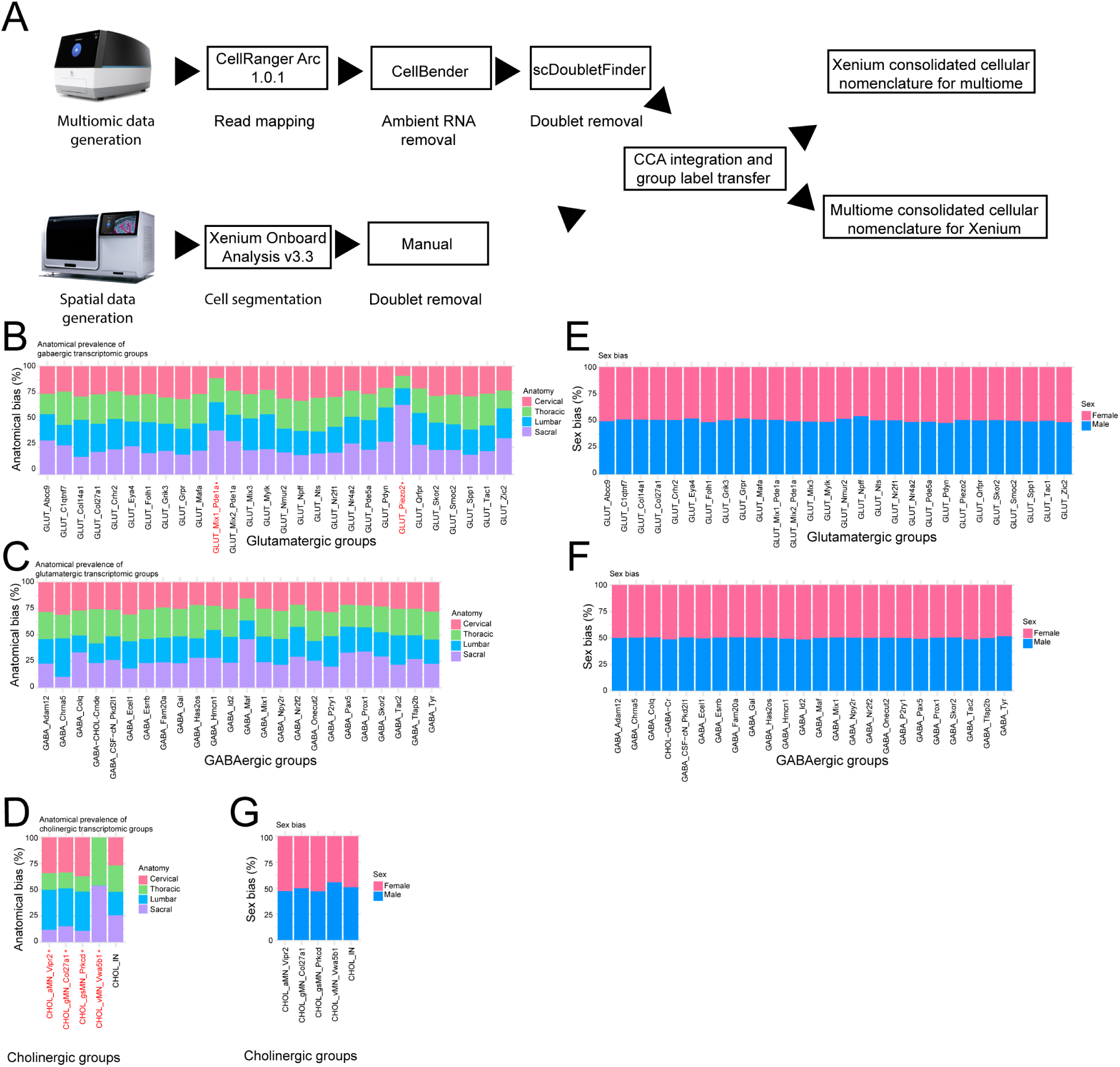
Aligning spatial and multiomic data and characterization of group level anatomical and sex biases in spatial transcriptomics profiled spinal cord data. (A) Workflow for integrating multiomic and spatial transcriptomic data (Chromium X and Xenium Analyzer images used with permission from 10x Genomics). (B) Rostro-caudal distribution of glutamatergic groups in a regionally balanced spatial transcriptomic sample (n= 270 844 cells). (C) Rostro-caudal distribution of GABAergic groups in a regionally balanced spatial transcriptomic sample (n= 228 784 cells). (D) Rostro-caudal distribution of cholinergic groups in a regionally balanced spatial transcriptomic sample (n= 17 616 cells). Groups marked by red asterisk/text are rostro-caudally significantly specialized (three-fold enrichment between at least two rostro-caudal regions, adjusted p<0.05 DeSeq2 and adjusted p<0.05 with beta regression). (E) Quantification of group level sex biases in sex balanced samples of Xenium profiled glutamatergic neurons (n = 312 660 cells, ns. for all groups as evaluated by DeSeq2, comparing models including and excluding categorical sex variable). (F) Quantification of group level sex biases in sex balanced samples of Xenium profiled GABAergic neurons (n = 270 626 cells, ns. for all groups as evaluated by DeSeq2, comparing models including and excluding categorical sex variable). (G) Quantification of group level sex biases in sex balanced samples of Xenium profiled cholinergic neurons (n = 24 384 cells, ns. for all groups as evaluated by DeSeq2, comparing models including and excluding categorical sex variable).

**Extended Data Figure 4:**
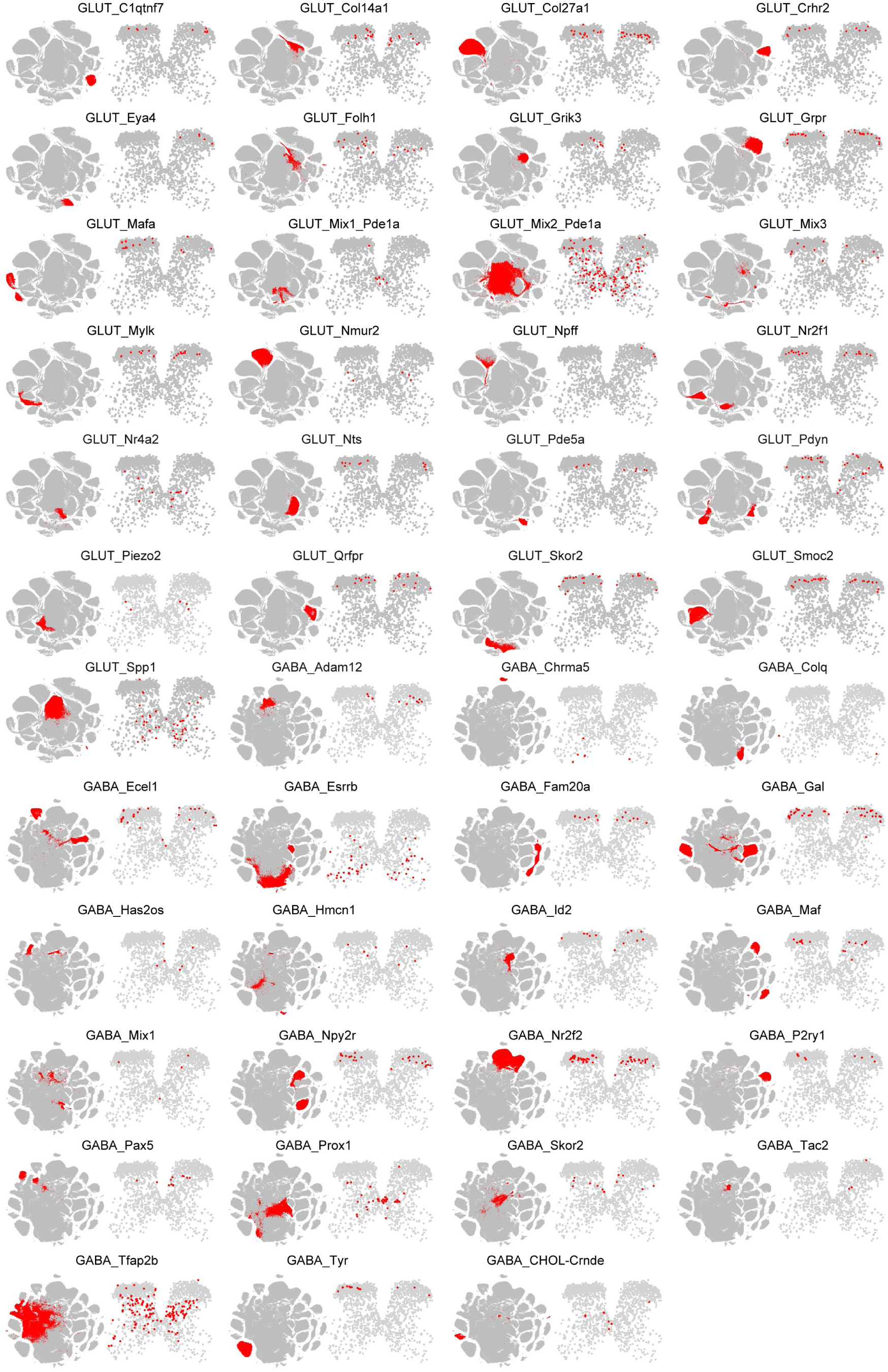
Group level anatomy for glutamatergic and GABAergic neurons. Groups highlighted in red in respective tSNE embeddings (left, glutamatergic or GABAergic cells) and corresponding anatomical distribution in a 10 micron lumbar spinal cord section (red highlight).

**Extended Data Figure 5:**
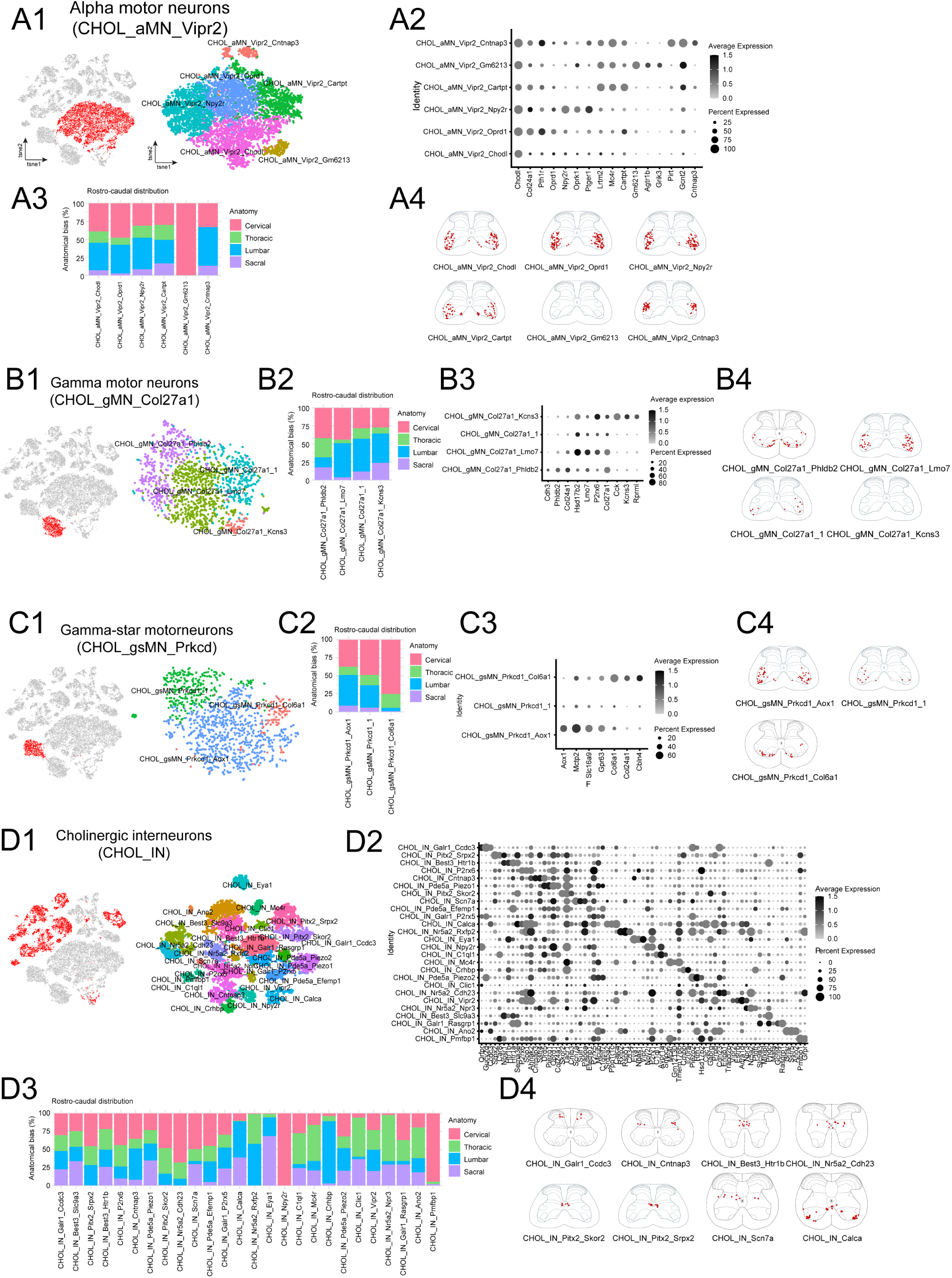
Transcriptomic diversity of the spinal cord cholinergic neurons profiled with Xenium spatial transcriptomics. (A1-A4) Transcriptomic diversity, molecular markers, rostro-caudal distribution and neuron type anatomy of the alpha motor neuron (CHOL_aMN_Vipr2) group of cholinergic neurons (n= 9008 cells). (B1-B4) Transcriptomic diversity, molecular markers, rostro-caudal distribution and neuron type anatomy of the gamma motor neuron (CHOL_gMN_Col27a1) group of cholinergic neurons(n= 1573 cells). (C1-C4) Transcriptomic diversity, molecular markers, rostro-caudal distribution and neuron type anatomy of the gamma-star motor neuron (CHOL_gsMN_Prkcd) group of cholinergic neurons (n= 1364 cells). (D1-D4) Transcriptomic diversity, molecular markers, rostro-caudal distribution and neuron type anatomy of the cholinergic interneuron (CHOL_IN) group of cholinergic neurons (n= 8498 cells).

**Extended Data Figure 6:**
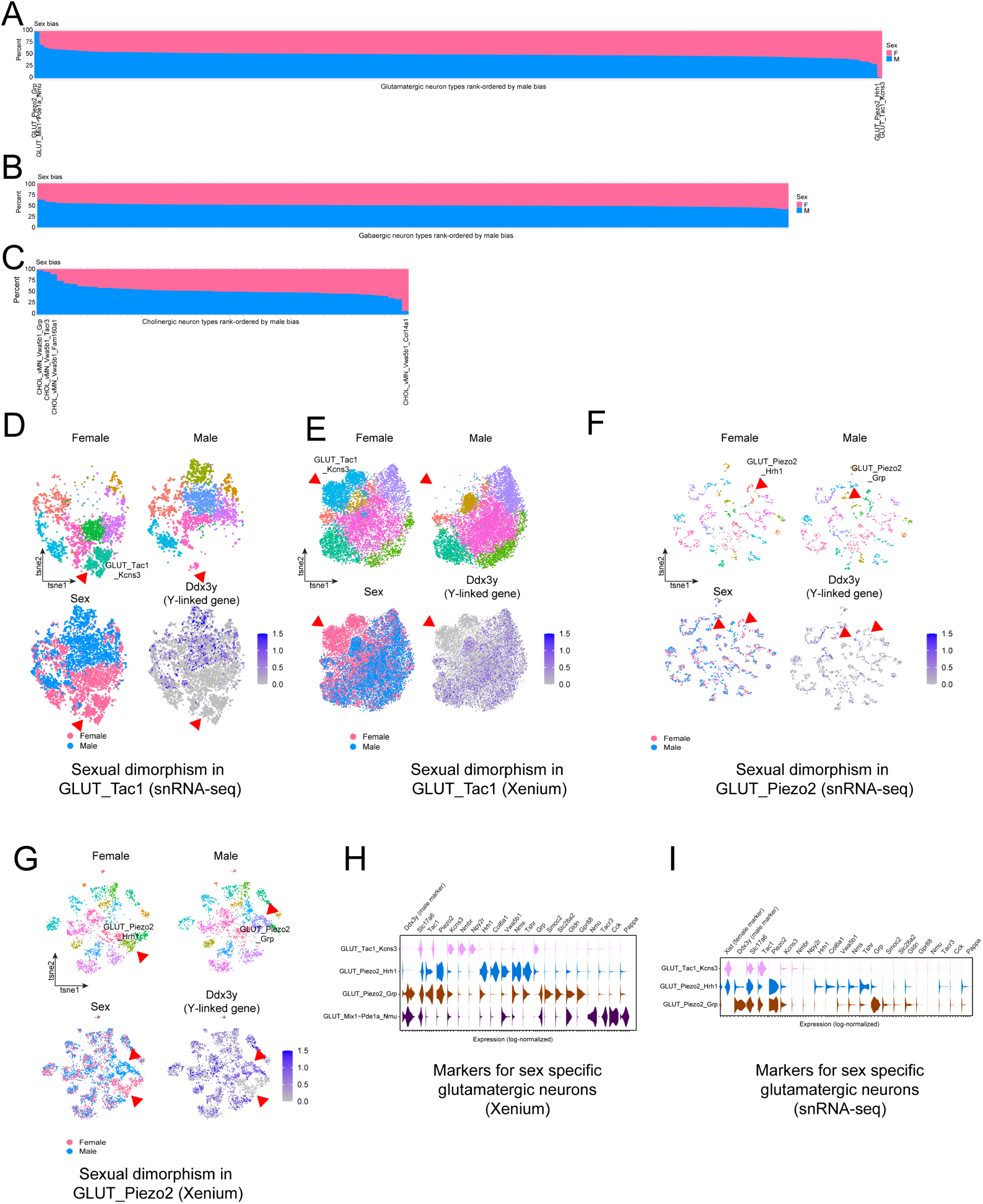
Analysis of sex specific neuron types in the adult spinal cord. (A) Rank-ordered glutamatergic neuron types based on male/female bias from sex balanced sample (n=14 mice per sex). (B) Rank-ordered GABAergic neuron types based on male/female bias from sex balanced sample (n=14 mice per sex). (C) Rank-ordered cholinergic neuron types based on male/female bias from sex balanced sample (n=14 mice per sex). (D) Sex specific neuron type in the multiome GLUT_Tac1 transcriptomic group. tSNE embedding of GLUT_Tac1 group neurons showing female (up left), and male (up right) cells, sex identity of neurons (lower left) and feature plot of the expression of the Y chromosome gene *Ddx3y* male marker (lower right, n= 6513 cells). Red triangles indicate sex specific glutamatergic neuron types. (E) Sex specific neuron type in the Xenium GLUT_Tac1 transcriptomic group. tSNE embedding of GLUT_Tac1 group neurons showing female (up left), and male (up right) cells, sex identity of neurons (lower left) and feature plot of the Y chromosome gene *Ddx3y* male marker gene expression (lower right, n= 20 272 cells). (F) Sex specific neuron type in the multiome GLUT_Piezo2 transcriptomic group. tSNE embedding of GLUT_Piezo2 group neurons showing female (up left), and male (up right) cells, sex identity of neurons (lower left) and feature plot of the expression of the Y chromosome gene *Ddx3y* male marker (lower right, n= 2381cells). (G) Sex specific neuron types in the Xenium GLUT_Piezo2 transcriptomic group. tSNE embedding of GLUT_Piezo2 group neurons showing female (up left), and male (up right) cells, sex identity of neurons (lower left) and feature plot of the Y chromosome gene *Ddx3y* male marker gene expression (lower right, n= 5438 cells). (H) Violin plot of marker genes defining sex specific glutamatergic neurons in the multiomic dataset. (I) Violin plot of marker genes defining sex specific glutamatergic neurons in the spatial transriptomic dataset.

**Extended Data Figure 7:**
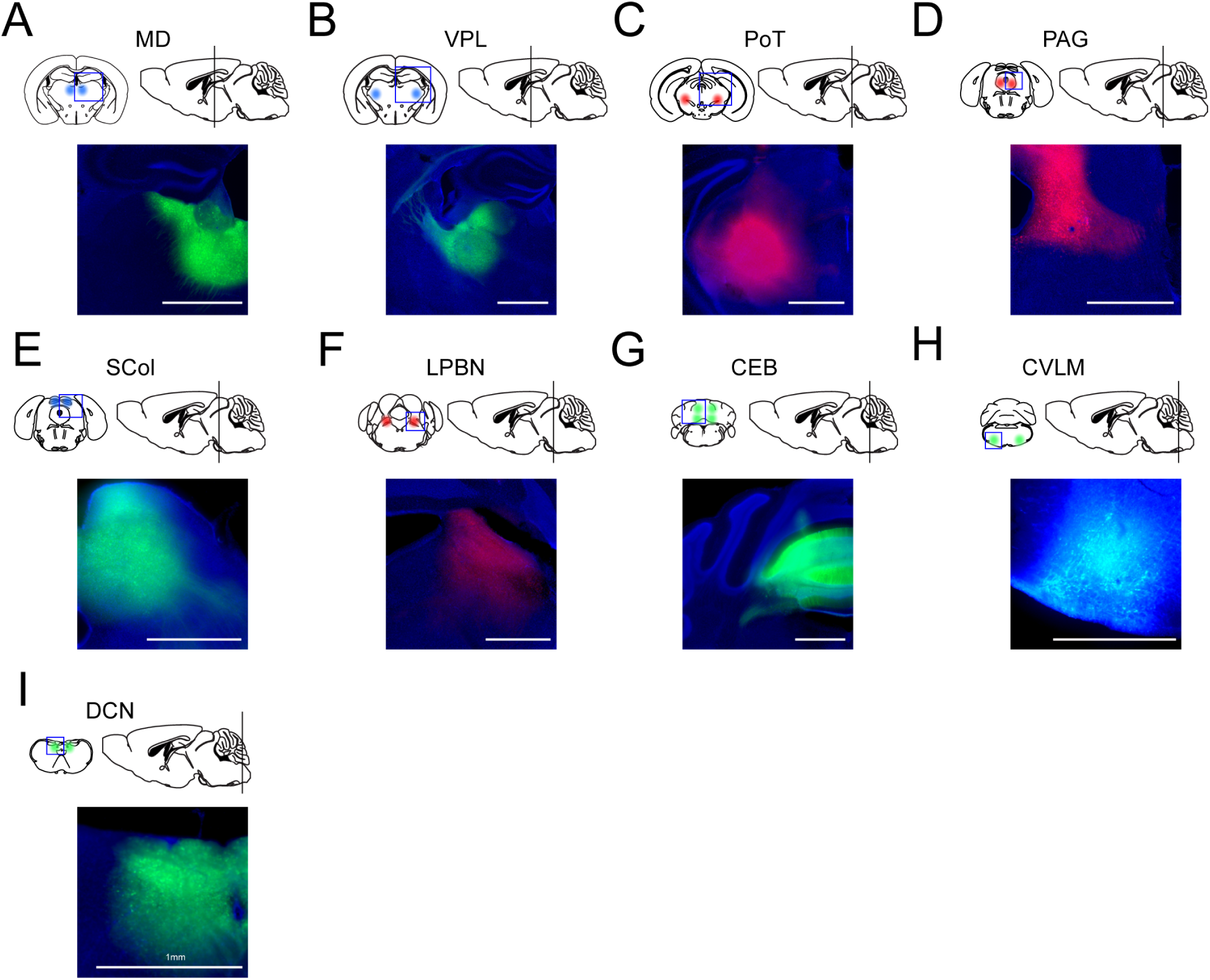
Retrograde tracing injection site validation. Epifluorescence images of retrograde tracer injection sites from: (A) MD (mNeonGreen); (B) VPL, (mNeonGreen); (C) PoT (tdTomato); (D) SCOL (mNeonGreen); (E) PAG (tdTomato); (F) LPBN (tdTomato). (G) CEB, (GFP); (H) CVLM, (GFP); (I) DCN (GFP). Blue channel – DAPI. Scale bar = 1 mm.

**Extended Data Figure 8:**
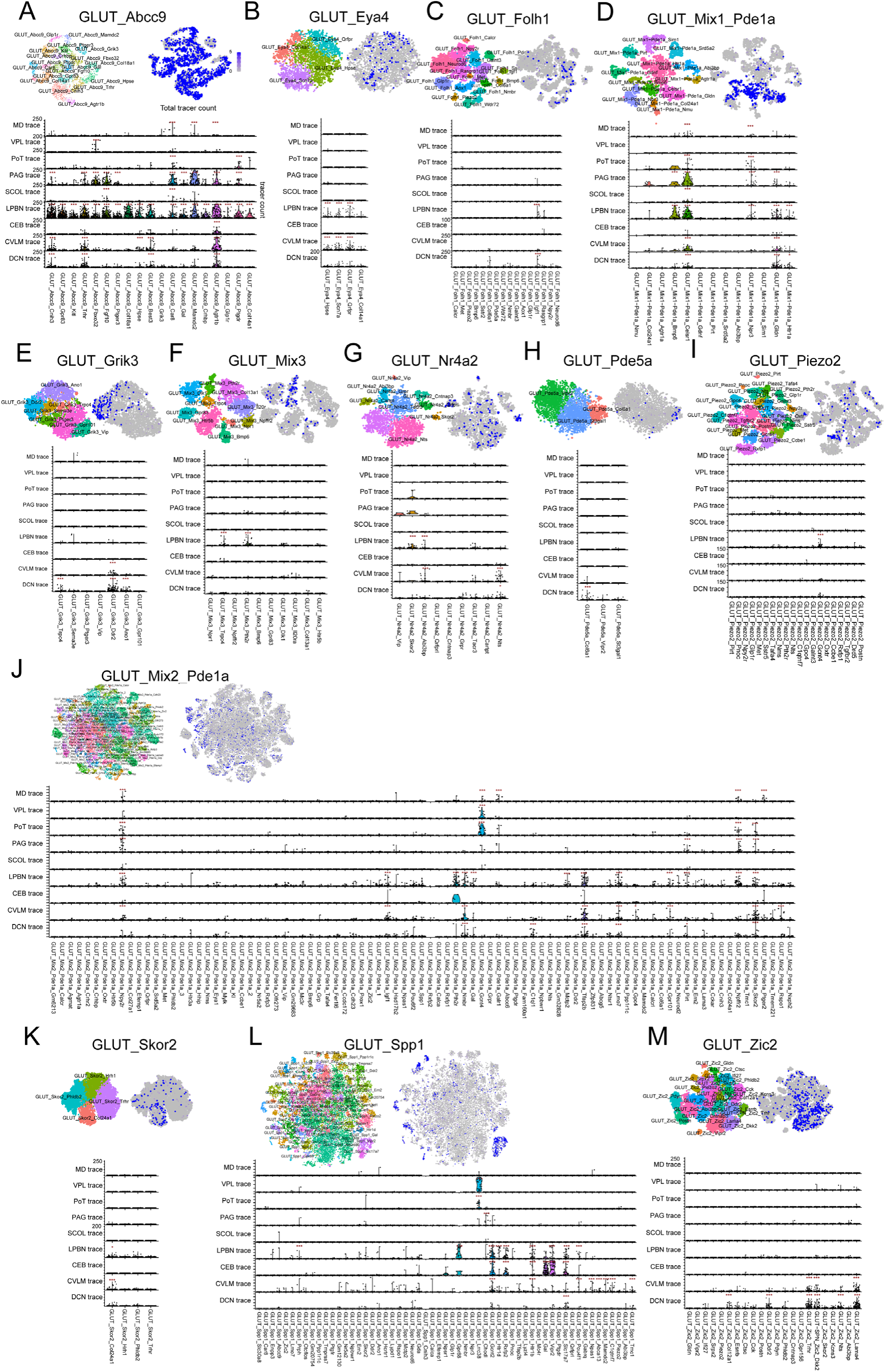
Spatial transcriptomics based identification of spinal projection neuron types by retrograde viral tracing. tSNE embedding of neuron types by group (up left), feature plot of total tracer count (up right) and violin plots of neuron type level tracer counts by traced structure (below) for groups containing significantly back-labeled neuron types including (A) GLUT_Abcc9, (B) GLUT_Eya4, (C) GLUT_Folh1, (D) GLUT_Mix1_Pde1a, (E) GLUT_Grik3, (F) GLUT_Mix3, (G) GLUT_Nr4a2 (H) GLUT_Piezo2, (I) GLUT_Pde5a, (J) GLUT_Mix2_Pde1a, (K) Glut_Skor2, (L) GLUT_Spp1, (M) Glut_Zic2 (n=4 animals per traced structure, permutation test with Bonferroni adjusted p values: * p<0.05, ** p<0.01, *** p<0.001).

**Extended Data Figure 9:**
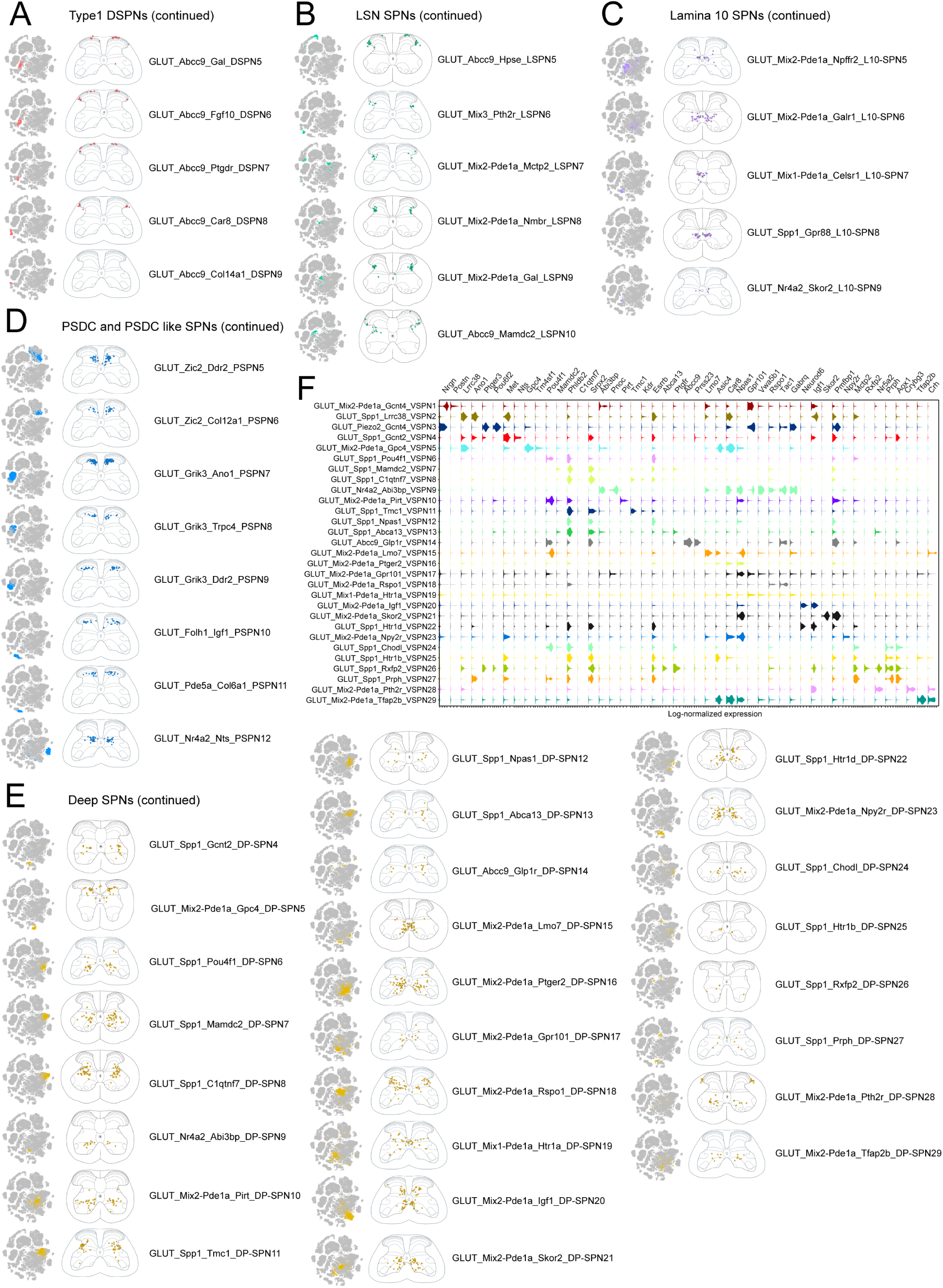
Molecular and anatomic catalog of mouse ascending spinal projection neurons (continued). (A) Anatomy for Type1 Dorsal SPNs. Continued from Figure 5C. (B) Anatomy for LSN SPNs (LSPNs). Continued from Figure 5E. (C) Anatomy for lamina 10 SPNs (L10-SPNs). Continued from Figure 5F. (D) Anatomy for postsynaptic dorsal column (PSDC) and PSDC-like SPNs (PSPNs). Continued from Figure 5H. (E) Anatomy for deep SPNs (DP-SPNs). Continued from Figure 5I. (F) Molecular markers (violin plot) for deep SPNs (DP-SPNs). For (A)-(E) cell type anatomical location was registered from 20 spinal cord 10 µm sections to reference lumbar (L5) spinal cord atlas or other rostro-caudal levels (cervical, thoracic, sacral) if cell type was not present in lumbar spinal cord.

**Extended Data Figure 10:**
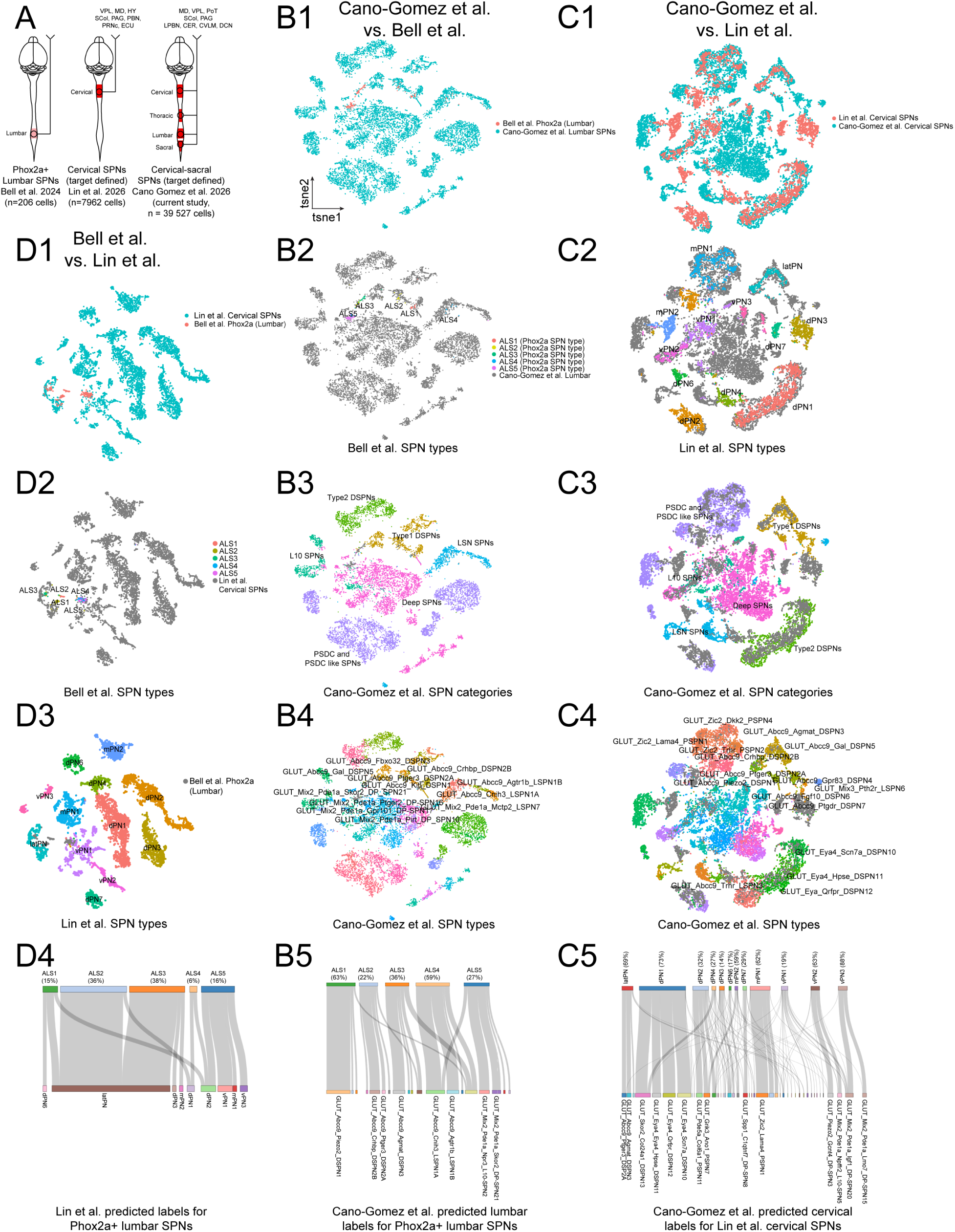
Alignment of spinal projection neuron atlases. (A). Comparison of analyzed spinal projection neuron single-cell transcriptomic profiling datasets. (B1) tSNE embedding of CCA aligned lumbar Phox2a+ projection neurons from Bell et al. 2024 (red, n = 206 cells) with lumbar SPN dataset from Cano-Gomez et al. (green, n=11 462 cells). (B2) Same as B1 with color coded lumbar Phox2a SPN types from Bell et al. 2024. (B3) Same as B1 with color coded lumbar SPN categories from Cano-Gomez et al. (B4) Same as B1 with selected color coded lumbar SPN types from Cano-Gomez et al. (B5) Sankey plot displaying corresponding neuron types identified by CCA label transfer from Cano-Gomez nomenclature to Phox2a+ nomenclature. Number in parenthesis reports the percentage of cells with high label transfer probability (>40%). (C1) tSNE embedding of CCA aligned cervical SPNs from Lin et al. 2026 (red, n = 7962 cells) with cervical SPN dataset from Cano-Gomez et al. (green, n=12 535 cells). (C2) Same as C1 with color coded cervical SPN types from Lin et al. 2026. (C3) Same as C1 with color coded cervical SPN categories from Cano-Gomez et al. (C4) Same as C1 with selected color coded cervical SPN types from Cano-Gomez et al. (C5) Sankey plot displaying corresponding neuron types identified by CCA label transfer from cervical Cano-Gomez et al. SPN dataset to Lin et al. cervical dataset. (D1) tSNE embedding of CCA aligned lumbar Phox2a+ projection neurons from Bell et al. 2024 (red, n = 206 cells) with cervical SPN dataset from Lin et al. 2026 (green, n = 7962 cells). (D2) Same as D1 with color coded lumbar Phox2a+ SPN types from Bell et al. 2024. (D3) Same as D1 with selected color coded cervical SPN types from Lin et al. 2026. (D4) Sankey plot displaying corresponding neuron types identified by CCA label transfer from Lin et al. 2026 to lumbar Phox2a lineage SPNs from Bell et al. 2024.

**Extended Data Figure 11:**
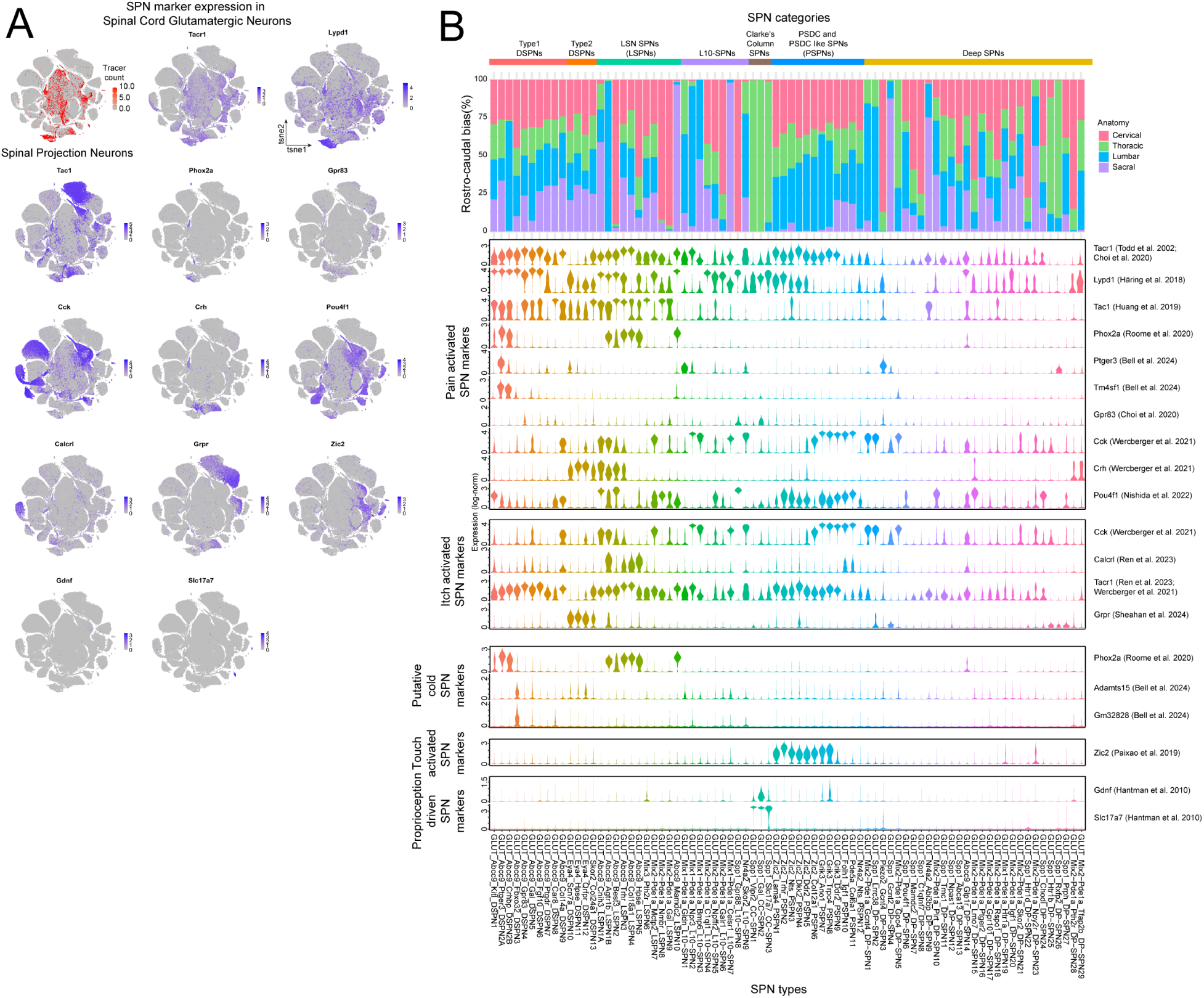
Specificity and expression pattern of previously identified SPN marker genes in the newly characterized SPN atlas. (A) SPN marker gene expression Feature plots in spinal cord glutamatergic neurons. Color scale: Loge-transformed expression. (B) Violin plots of SPN marker genes in the SPN atlas (below, log_e_ transformed expression). Rostro-caudal bias and anatomical category of individual SPN types (up).

